# A method to enrich polypeptidyl-tRNAs to capture snapshots of translation in the cell

**DOI:** 10.1101/2022.04.22.489242

**Authors:** Ayako Yamakawa, Tatsuya Niwa, Yuhei Chadani, Akinao Kobo, Hideki Taguchi

## Abstract

Life depends on proteins, which all exist in nascent states when the growing polypeptide chain is covalently attached to a tRNA within the ribosome. Although the nascent chains; *i*.*e*., polypeptidyl-tRNAs (pep-tRNAs), are considered as merely transient intermediates during protein synthesis, recent advances have revealed that they are directly involved in a variety of cell functions, such as gene expression control. An increasing appreciation for fine-tuning at translational levels demands a general method to handle the pep-tRNAs on a large scale. Here, we developed a method termed *pe*ptidyl-*t*RNA *e*nrichment using *o*rganic extraction and *s*ilica adsorption (PETEOS), and then identify their polypeptide moieties by mass spectrometry. As a proof-of-concept experiment using *Escherichia coli*, we identified ∼800 proteins derived from the pep-tRNAs, which were markedly biased towards the N-termini in the proteins, reflecting that PETEOS captured the intermediate pep-tRNA population during translation. Furthermore, we observed the changes in the pep-tRNA set in response to heat shock or antibiotic treatments. In summary, PETEOS will complement conventional methods for profiling nascent chains such as ribosome profiling.

**Significance Statement:** In the central dogma of biology, RNA and protein are usually regarded as two completely independent molecular species. However, they are combined into a single species called peptidyl-tRNA (pep-tRNA) during the translation process in the ribosome. Despite the importance of pep-tRNAs as precursors of all proteins in the cell, a general method to analyze pep-tRNAs on a large scale was lacking. Taking advantage of the properties of pep-tRNAs as RNA and protein, we developed a method to enrich the pep-tRNAs by organic solvent extraction and silica column separation. The method, termed PETEOS, not only provides a unique approach to examine the nascent state of proteins but also may be effective in capturing snapshots of translation status in the cell.

## Introduction

Every protein in the cell is generated through translation. The formation of the proteome involves the regulation of various steps in gene expression. In addition to the well-known importance of gene regulation at transcriptional levels, gene expression control at translational levels has been increasingly appreciated (1, 2).

The translational control is diverse, flexible, and rapid, depending on cellular contexts or stimuli such as a variety of stresses. In addition to the control at the translation initiation step, which is a major rate-determining step in translation, even the translation elongation could be regulated (3, 4). One of the fine-tuning events is elongation rate control, as exemplified by elongation pausing, which can affect the folding, function, and quality control of proteins (3–7). For example, the translation product of the gene *secM* in *Escherichia coli* induces translation arrest (8, 9). The translation arrest by SecM is directly involved in the feedback regulation of its downstream gene, *secA* (8, 9). In addition to the ad hoc identification of translational pausing, biochemical analyses of over 1,000 *E. coli* genes revealed that a majority of them indeed undergo single or multiple pauses in translation, indicating the widespread occurrence of translational pausing (10). Since it is evident that not only pausing but also many other biological events are regulated at the translational level, there is an increasing demand for methods to globally investigate the translation process in the cell.

So far, two main methodologies based on completely different concepts are used to survey the translation process in cells (11). One is ribosome profiling, a deep sequencing technique for ribosome-protected mRNA fragments generated by RNase digestion (12). Ribosome profiling is a very powerful genome-wide analysis with nucleotide resolution and has been established as a method to monitor the translation status in cells (11–16). However, this method, which relies on a huge amount of “ribosome footprint” sequencing data, is inherently indirect in detecting the nascent proteins or pep-tRNAs, and might cause some controversy about how to collect and interpret the data (*e*.*g*. 17). In addition, the method is still time-consuming and costly.

The other methodology is based on proteomics with liquid chromatography-tandem mass spectrometry (LC-MS/MS). Several “translatome” methods have been developed to distinguish nascent proteins from the pre-existing protein population. The proteomics-based approaches rely on the labeling of nascent polypeptide chains, using noncanonical amino acids such as azide homoalanine (*e*.*g*., BONCAT (18), QuaNCAT (19)) and puromycin or its analogs (*e*.*g*., PUNCH-P (20, 21), OPP-ID (22), pSNAP (23)). Since the vast majority of nascent proteins labeled by the noncanonical amino acids have already been released from the ribosome, these proteins are newly but already “synthesized” mature proteins, and not nascent proteins during translation. The puromycin-related methods rely on the principle that puromycin or its analogs terminate translation by entering the A-site of the ribosome, and are then covalently incorporated into the growing end of the nascent peptide moiety in pep-tRNAs. However, certain nascent peptides are resistant to puromycin (24–26). In addition, all of these methods that rely on labeling usually require an hour-long timescale or more to accumulate the labeled products (27). In summary, these analytical methods for the nascent proteome do not directly deal with pep-tRNAs and do not capture the moments of the translation elongation status in the cell.

In view of these points, a direct methodology to handle pep-tRNAs is required to complement the prevailing methods for the translatome analyses. Although the point is not always appreciated, it is obvious that all proteins are born via pep-tRNAs, which possess the properties of both RNA and protein. In other words, pep-tRNA is an essential translation intermediate that lies between RNA and protein in the central dogma of biology. Despite their importance in life, pep-tRNAs are low-abundant. Assuming that the approximate numbers of proteins and ribosomes in *E. coli* are ∼3 million and ∼70,000, respectively (28), the maximum amount of pep-tRNAs is at most only ∼2% of total proteins, even if all ribosomes participate in translation.

These “in-between” properties of both RNA and protein and the low abundance make pep-tRNAs challenging to handle on the “pep-tRNA-ome” scale. Ito *et al*. developed an ingenious method to collectively detect cellular pep-tRNAs, the “nascentome,” using a combination of pulse-labeling and two-dimensional electrophoresis (29). The “nascentome” analysis can detect the whole pep-tRNAs but cannot identify which protein is part of each pep-tRNA. Therefore, we developed a non-labeling method to enrich and identify pep-tRNAs in cells. Briefly, pep-tRNAs in cells are enriched by organic solvent extraction, followed by isolation on a silica column. The polypeptide moieties in the enriched pep-tRNAs are cleaved under high pH and high temperature conditions. The resultant polypeptides are digested by proteases such as trypsin and then identified using conventional LC-MS/MS-based shotgun proteomics. This method, termed *pe*ptidyl-*t*RNA *e*nrichment using *o*rganic extraction and *s*ilica adsorption (PETEOS), is not only complementary to prevailing translation measurement methods, but also has, in principle, several advantages such as rapid capturing of translation status and non-labeling feature.

## Results

### Polypeptidyl-tRNA enrichment and identification using a proteomics approach

As far as we know, there is no established method to purify whole pep-tRNAs from cells. Our strategy to enrich pep-tRNAs takes advantage of their two distinctive properties as RNA and polypeptide. The method, which we call PETEOS, is composed of four steps (Fig. 1A). Step 1: Enrichment of pep-tRNAs and proteins from cells. Harvested cells are directly mixed into a phenol-containing reagent (*e*.*g*. TriPURE, TRIzol) to separate most RNAs including ribosomal RNAs (aqueous phase) from pep-tRNAs and proteins (organic phase). Note that there is no need for labeling times in this method. After the chloroform addition, the pep-tRNAs and proteins in the organic phase and interphase are precipitated by trichloroacetic acid (TCA)/acetone, under conditions where the ester bonds between polypeptide and tRNA in the pep-tRNAs are retained. Step 2: Isolation of the pep-tRNAs. The RNA is isolated by using silica to adsorb the nucleic acid portions of pep-tRNAs, under highly concentrated chaotropic salt conditions (30, 31). Step 3: Cleavage of peptide moieties from peptidyl-tRNAs. The ester bond between peptide and tRNA is cleaved by high pH and heat treatment (29). Step 4: Identification of the peptides after trypsin treatment, by LC-MS/MS-based shotgun proteomics.

**Figure 1.**
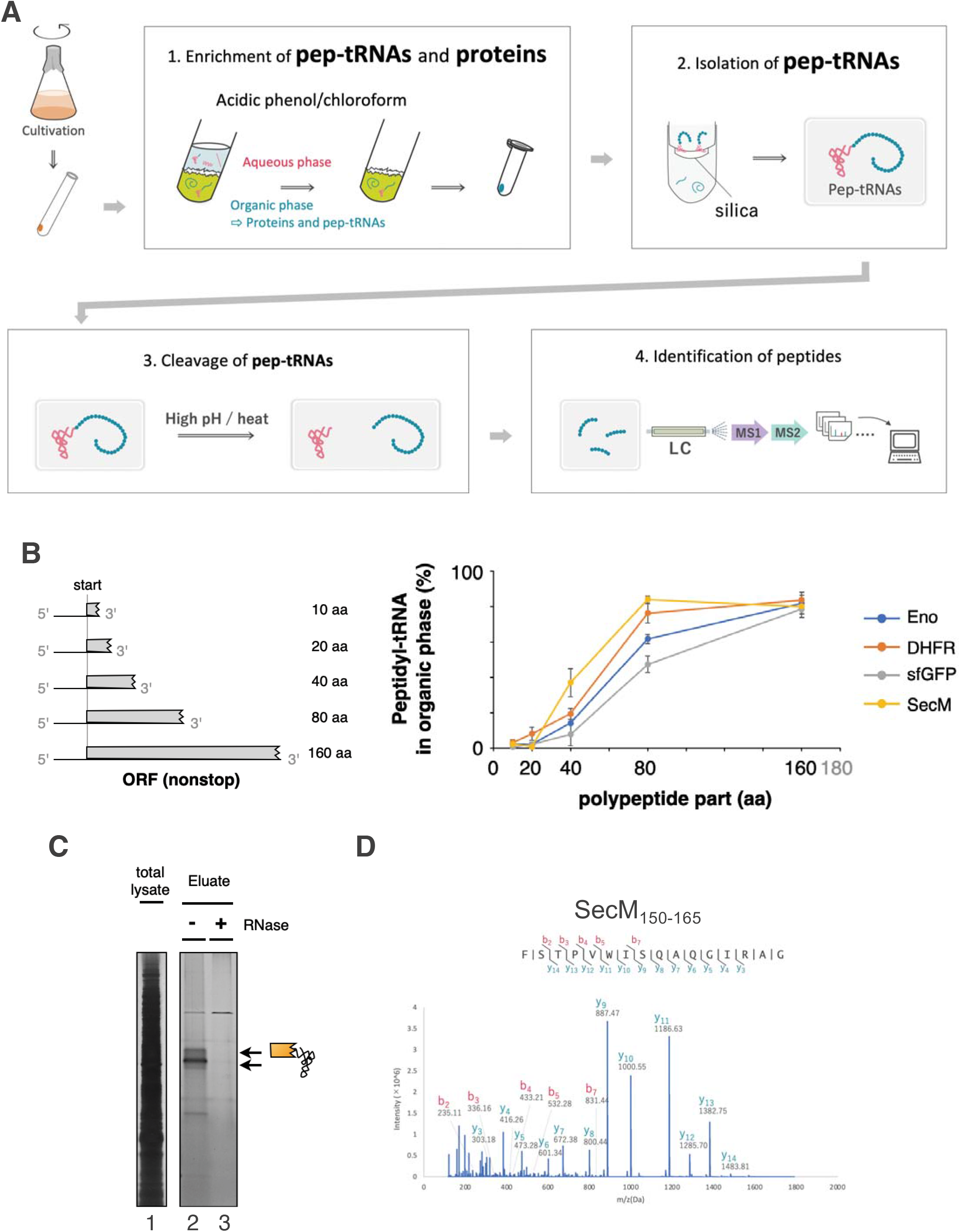
PETEOS workflow to enrich and identify polypeptidyl-tRNAs (pep-tRNAs) in cells. **A. Step 1**: Enrichment of pep-tRNAs and proteins. Log phase *E. coli* cells (OD660 ∼ 0.5) cultured at 37 °C are extracted by phenol/chloroform to capture the hydrophilic RNA molecules into the aqueous phase. Relatively hydrophobic proteins and pep-tRNAs in the organic phase are precipitated by TCA/acetone. **Step 2**: Isolation of the pep-tRNAs by a silica column in the presence of concentrated chaotropic salts. **Step 3**: Hydrolysis of the ester bond between peptide and tRNA in pep-tRNA under high pH and high temperature. **Step 4**: Identification of the peptide moieties of the pep-tRNAs by conventional shotgun proteomics after trypsin digestion. **B**. Fractionation of pep-tRNAs by phenol-extraction. *Left*, schematic of the truncated nonstop mRNAs to prepare the pep-tRNA with various lengths of the polypeptide moiety. *Right*, the fractionation ratio of the pep-tRNA into the organic phase was calculated from the gel image (representative in Fig. S1B) as described in materials and methods. Pep-tRNAs carrying N-terminal 10, 20, 40, 80 or 160 aa of *E. coli* enolase (Eno), DHFR, *secM*, or superfolder GFP mRNAs were examined. The mean ± SE values of three independent experiments are shown. **B**. Purification of SecM-tRNA by silica column. The lysate of *E. coli* cells overexpressing *secM* was processed by silica column purification as described in materials and methods. Cellular extract (lane 1) and the eluate with (lane 3) or without (lane 2) RNase treatment during the isolation procedure were separated by neutral pH SDS-PAGE and developed by silver staining. **C**. MS2 spectra of the C-terminal peptides of SecM (_150_FSTPVWISQAQGIRAG_165_) from the *E. coli* cells overexpressed SecM.

### Validation of the phenol extraction (Step 1)

The validity of the PETEOS method using *E. coli* lysates was evaluated at each step. An LC-MS/MS analysis of the TCA/acetone precipitate in Step 1 (organic phase) revealed that ∼79% of the identified proteins in the precipitate overlapped with proteins identified by a conventional method (32) to extract proteins from cells (Fig. S1A), confirming that the protein fraction in the organic phase seems to lack significant bias. Even if there was no bias in the total protein fraction, we need to know how peptidyl-tRNAs are partitioned in organic solvent extraction in Step 1 since the chemical properties of pep-tRNAs with various polypeptide moieties are diverse. For example, when the peptide moiety is hydrophilic or relatively short, the hydrophilic property of the RNA would be dominant, but when the polypeptide portion is more hydrophobic or longer, the more protein-like property would be manifested.

Then, we investigated this issue in a model system. First, we focused on the length of the peptide moieties of pep-tRNAs. To prepare the pep-tRNA with various peptide lengths, we translated nonstop mRNAs in a reconstituted *in vitro* translation system (33). The 3’ end of *E. coli* enolase (Eno), DHFR, *secM*, or superfolder GFP mRNAs were truncated at 30, 60, 120, 240, or 480 nucleotides (10, 20, 40, 80, or 160 amino acids) from the start codon, respectively (Fig. 1B). The reaction mixture was then phenol-extracted. The pep-tRNA in each fraction was analyzed by SDS-PAGE under neutral pH conditions (neutral-PAGE), where the ester bonds between peptide and tRNA in the pep-tRNAs remain intact, as described previously (34) (Fig. S1B). As shown in Fig. 1B, all the pep-tRNAs we tested were fractionated into the organic phase in a length-dependent manner. Furthermore, the hydrophobicity of the polypeptide moiety, though not as much as its length, was shown to have some influence on the phenol-based fractionation (Fig. S1C). These results provided us with some tendencies in the distribution of pep-tRNA in the phenol extraction, where pep-tRNAs with relatively short peptide moieties do not distribute well in the organic phase.

### Validation of the later steps (Step 2-4) of the PETEOS method

Next, we analyzed the silica-based concentration of the pep-tRNA from the cell extract (Step 2). Accordingly, we overexpressed a well-known pausing peptide, SecM. Then we analyzed the eluate of silica-isolation by SDS-PAGE and following silver staining. SecM pep-tRNA was successfully isolated from the cellular extract (Fig. 1C). Furthermore, the MS/MS analysis identified the SecM-derived peptides including the _150_FSTPVWISQAQGIRAG_165_ fragment from this purified fraction (Fig. 1D). The translation of the gene *secM* is arrested at the 165^th^ glycine, where tRNA^Gly^ is covalently attached to the elongating end of the nascent peptide (SecM_1-165_-tRNA^Gly^). This suggests that the PETEOS method has a potential to identify paused peptides.

A previous nascentome study showed that Tris-base in the buffer stimulates the ester bond cleavage of pep-tRNAs (29). A nucleophilic attack of tris(hydroxymethyl)aminomethane (Tris) on the carboxyl group of the C-terminal amino acid, which is ester-bonded to tRNA in the pep-tRNA, produces a Tris-adduct during the pep-tRNA hydrolysis (Fig. S1D). Therefore, we speculated that some peptides derived from pep-tRNAs could become conjugated with Tris. A peptide search assuming the Tris-addition at the C-terminus revealed ∼10 Tris-conjugated peptides, including a SecM-derived paused peptide (Fig. S1E). The conjugation of the Tris-base confirmed the handling of intact pep-tRNAs until they are exposed to high pH and high temperature.

### Proof-of-concept analysis using *Escherichia coli* cell lysates

The silica-eluate isolated from the organic phase including the components of exponentially growing *E. coli* cells (at Step 2) was separated by neutral-PAGE analysis. We observed smeared silver-stained bands (Fig. 2A). Since an RNase treatment before the adsorption to the silica column drastically reduced the smeared bands (Fig. 2A), they would be derived from pep-tRNA species.

**Figure 2.**
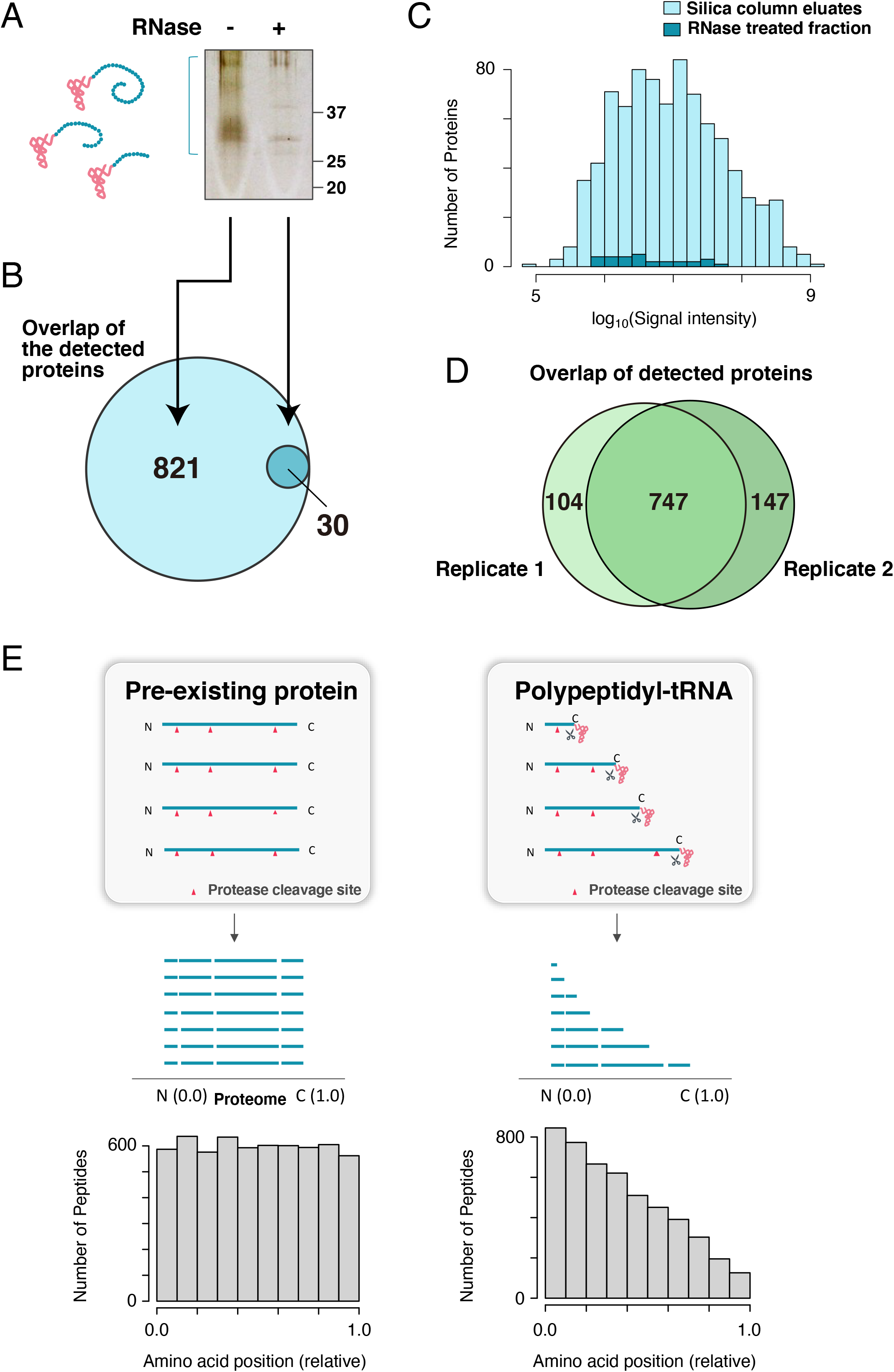
Enrichment and identification of pep-tRNAs in *E. coli*. **A**. The eluate from the silica column (Step 2) was analyzed by SDS-PAGE under neutral pH, and detected by silver staining. The eluate with an RNase treatment after Step 1 was also analyzed. Pep-tRNA molecules are schematically indicated (tRNA moiety, orange; peptide moiety, blue). **B**. Number of proteins identified as pep-tRNA molecules in the method. The proteins with < 1% FDR and ≥ 2 number of detected peptides were counted. All 30 proteins identified in the RNase-treated fraction were also detected from the no-treatment fraction. **C**. Histogram of summed peptide intensity per protein (log10 scale) obtained by MS1 peaks for proteins from the silica column eluate or the RNase-treated fraction. **D**. Overlap of the detected proteins between two replicates. The proteins with < 1% FDR and ≥ 2 number of detected peptides were counted. **E** (Upper) Illustration of pep-tRNAs and pre-existing proteins digested by trypsin. Trypsin digestion of pep-tRNAs, which elongate from the N- to the C-terminus, should accumulate more peptides containing the N-terminal portion. In contrast, for pre-existing proteins, which basically have full-length polypeptides, the amounts of trypsin-digested peptides should be equal for all peptides. Arrowheads colored in red indicate the sites for the trypsin cleavage. Relative positions of the N-terminus and C-terminus are defined as 0 and 1, respectively. (Lower) Meta-analysis of the relative position of the peptides identified by LC-MS/MS for pep-tRNAs and pre-existing proteins.

After the peptide moieties from the pep-tRNAs were obtained by the alkaline and heat treatment (Step 3), the LC-MS/MS analysis (Step 4) identified ∼5,000 peptides derived from ∼850 proteins (Fig. 2B, Fig. S2A, and Supplementary Dataset S1). The number of the detected peptides and proteins was drastically decreased in the RNase-treated fraction (Fig. 2B, Fig. S2A, and Supplementary Dataset S1), indicating that more than 95% of the identified proteins would be derived from pep-tRNAs. In addition, the signal intensities of the proteins derived from the RNase-treated fraction were much lower than those without the RNase treatment (Fig. 2C). Furthermore, the proteins detected in the RNase-treated fraction include many high abundant proteins such as chaperones, translation factors, and several outer membrane proteins, suggesting that these proteins were probably derived from non-specific interaction with the silica column (Supplementary Dataset S1). From these results, we concluded that the PETEOS method can enrich and identify the cellular pep-tRNAs.

We then investigated the overlap between the detected proteins by the PETEOS method and a conventional proteome method. The result showed that ∼80% of the identified proteins by the PETEOS method were also identified by a conventional method (Fig. S2B).

Furthermore, we examined the reproducibility of this method by comparing the results between two biological replicates. Among ∼850 detected proteins in each replicate, ∼80 % were overlapped (Fig. 2D). At the peptide level, ∼70 % of the peptides were overlapped (Fig. S2C). Quantitative comparison by MS1 intensities revealed that the abundance ratios between two replicates were within the 5-fold range (Fig. S2D). Although precise quantitative assessment seems difficult compared to conventional quantitative proteomics for whole-cell lysates (usually ∼2-fold changes are detectable), the PETEOS method has the potential to detect somewhat large proteomic changes.

We next investigated whether the MS-identified peptides derived from the pep-tRNAs in *E. coli* are in fact molecular species during the process of translation. Since nascent peptides elongate from the N- to C-terminus, we expected that the tryptic peptides derived from pep-tRNAs would be enriched with the N-terminus as compared to those derived from pre-existing total cellular proteins (Fig. 2E, upper panel). A meta-analysis demonstrated that the tryptic peptides from the pep-tRNAs were highly biased toward the N-terminal regions (Fig. 2E, lower panel). In contrast, a meta-analysis of the total pre-existing proteins showed equally distributed tryptic peptides (Fig. 2E, lower panel), as expected since the total proteins are basically composed of the full-length polypeptides after translation. The analysis confirmed that the MS-identified peptides derived from the pep-tRNAs in *E. coli* are indeed intermediate species during the translation elongation.

### Nascentome change in response to heat stress

To test whether the PETEOS method could capture the rearrangement of the nascentome upon environmental perturbations, we evaluated the nascentome change by heat treatment with the PETEOS method. The cells were cultivated at 45 °C for 30 min and disrupted with the phenol reagent. Then, pep-tRNAs were isolated, and the peptides moieties in the pep-tRNAs were analyzed. The peptides detected from the samples again showed quite high sensitivities to RNase treatment and positional biases toward the N-termini, confirming that the PETEOS method is robust to heat treatment (Fig. S3A). Quantitative assessment of the peptides obtained from the heat-treated and non-treated cells showed that several peptide fragments were specifically increased or decreased by the heat treatment (Fig. 3A and Supplementary Dataset S2). Among them, the peptides derived from heat shock proteins and proteases (GroEL/ES, DnaK, ClpB, HtpG, Lon, and FtsH) were strongly biased toward the enriched fraction (Fig. 3A and Supplementary Dataset S2). Importantly, the abundance ratios (fold changes) of these Hsps and proteases obtained by the PETEOS method were much larger than those by the conventional whole-cell proteome experiment (Fig. 3B and Supplementary Dataset S3). These results suggest that acute protein expression under the heat shock condition is strongly biased beyond the changes observed by the conventional proteome analysis. In other words, we can investigate such a “snapshot” of the translation status using the PETEOS method.

**Figure 3.**
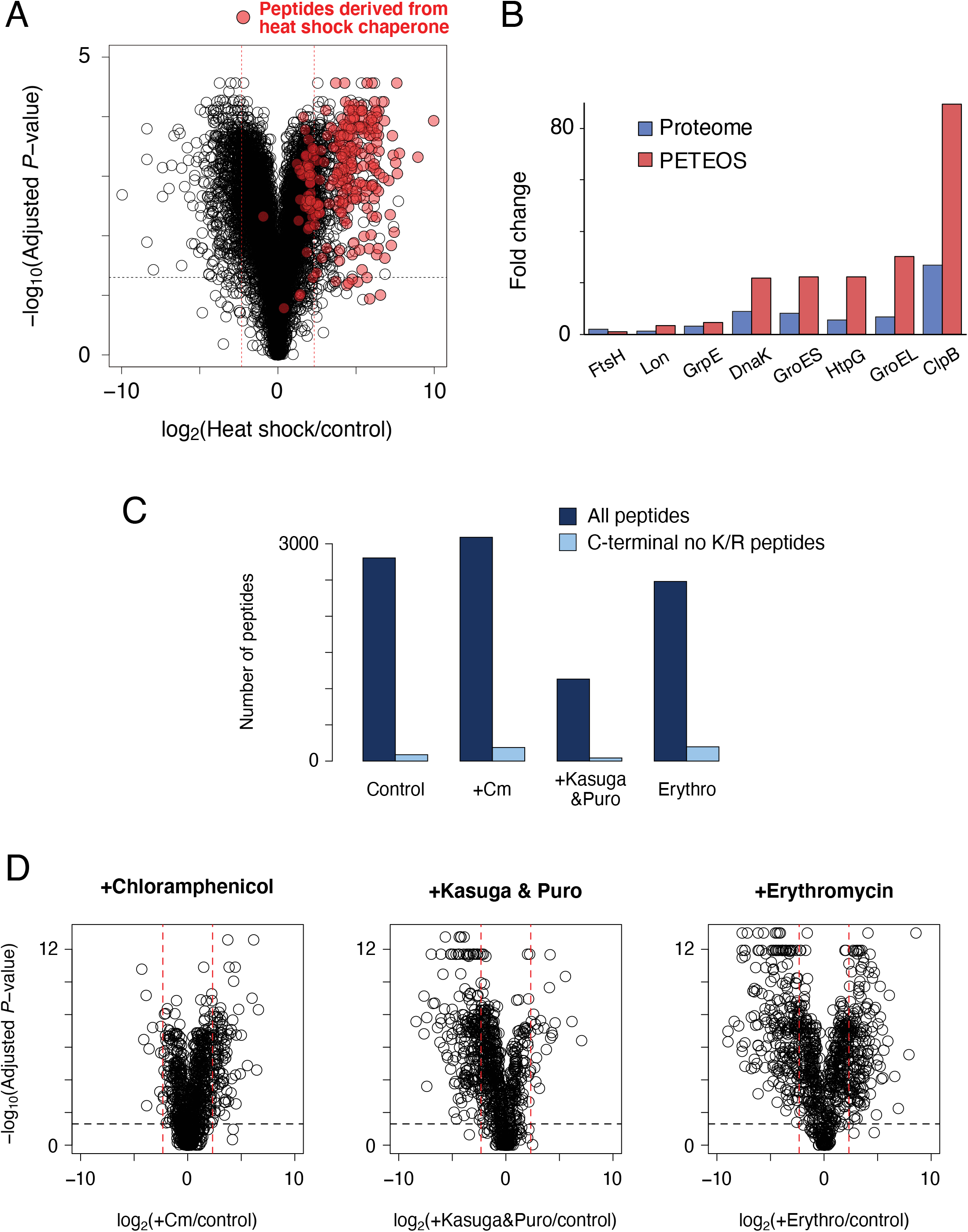
Investigation of the changes in the nascent peptides/proteins by heat-shock and antibiotics. **A**. Quantitative assessment of the changes by the heat-shock (45 ºC, 30 min) at the peptide level. Red dots indicate the peptides derived from the heat shock proteins and proteases (GroEL/ES, DnaK, ClpB, HtpG, Lon, and FtsH). Red dotted lines represent 5 and 1/5 fold change values, indicating around the 2*σ* point obtained by comparing two replicates (see Fig. S2D). The horizontal axis indicates the value of the fold changes by the heat shock taken as log_2_, and the vertical axis indicates *P*-values by t-test with three technical replicates in each sample taken as −log_10_. The values of the fold changes and *P*-values were obtained by Proteome Discoverer 2.4 (detailed settings were described in the Materials and Methods section and Supplementary Table S3). The peptides with less than three number of peptide-spectrum matches (PSMs) were omitted. **B**. Fold changes of the heat shock proteins evaluated by the PETEOS method and a conventional proteome method. The fold change data by the conventional method was acquired by the SWATH-MS acquisition method. **C**. Number of the detected peptides from the antibiotics-treated samples. The number of the peptides whose C-termini was not Lys (K) or Arg (R) were also shown. The peptides with less than three number of PSMs were omitted. **D**. Quantitative assessment of the changes by treating the antibiotics at the protein level. Red dotted lines represent 5 and 1/5 fold change values, indicating around the 2-sigma point obtained by comparing two replicates (see Fig. S2D). The horizontal axis indicates the value of the fold changes by the treatment taken as log_2_, and the vertical axis indicates *P*-values by t-test with three technical replicates in each sample taken as −log_10_. The values of the fold changes and *P*-values were obtained by Proteome Discoverer 2.4 (detailed settings were described in the Materials and Methods section and Supplementary Table S3).

### Nascentome change in response to antibiotic treatments

As another example of the application of the PETEOS method, we examined the rearrangement of the nascentome by the treatment of several antibiotics that inhibit translation reactions on ribosomes. For this purpose, we employed three conditions: chloramphenicol, kasugamycin+puromycin, and erythromycin treatment. Chloramphenicol inhibits the peptidyl-transferase reaction of ribosomes, so that the change in the snapshot of the protein synthesis is expected to be small, or the efficiency of pep-tRNA detection may be better (35). Puromycin accepts the polypeptide part from pep-tRNA in a ribosome function-dependent manner(36). Therefore, the pep-tRNAs that are detectable by PETEOS are anticipated to be decreased upon the puromycin treatment. In addition, the property of puromycin may allow us to assess the puromycin-resistant pep-tRNAs that impair ribosomal function to pause the translation elongation. For this purpose, we added the kasugamycin in parallel to highlight the effect of puromycin by inhibiting the following translation initiation. Erythromycin, a kind of macrolide antibiotic, binds to the exit tunnel of the ribosome to inhibit translation elongation. Previous studies unveiled that translation inhibition by the macrolide has some specificity for the nascent peptide sequence within the tunnel (37, 38).

The cells were treated with each antibiotic or cocktail of antibiotics for 10 min and disrupted with the phenol reagent. The identified peptide sets, which had properties as nascentome (Fig. S3B), showed antibiotic-dependent characteristics (Fig. 3C, D). As expected, chloramphenicol treatment caused a small increase in the number of detected peptides (Fig. 3C) and the abundance ratio of a subset of proteins (Fig. 3D). On the other hand, the kasugamycin/puromycin treatment caused a strong decrease in the number of the detected peptides (Fig. 3C) and fold changes of the many proteins (Fig. 3D). However, we observed several pep-tRNAs such as a fragment of SecM with unchanged abundances even after the treatment, suggesting that ribosomes accommodating these pep-tRNAs poorly react with puromycin.

The treatment by erythromycin resulted in both an increase and decrease in subsets of proteins. We then evaluated the relationship between the abundance changes and the sequence motif of erythromycin (“[K/R]-X-[K/R]”) reported previously (37, 38). The result showed the decreasing tendency of the abundances as the number of erythromycin-dependent pausing motifs increased (Fig. S3C). This bias could be caused by ribosome rescue reaction at the erythromycin-paused ribosome (39).

Finally, since the PETEOS method relies on the shotgun proteomics approach with tryptic digestion and electrospray ionization mass spectrometry, a large bias exists in the detectable peptides. Hence, it is usually difficult to discuss the precise position of translational pausing. However, we detected several peptides whose C-termini were not lysine or arginine, implying that these peptides are the C-terminal region of the pep-tRNAs since they were not considered to be products of the tryptic digestion (Supplementary Table S1). In addition, some of these peptides were also detected as C-terminal Tris-added peptides, confirming that they were in the C-terminal region of the pep-tRNAs. Although the number of these peptides was not large, these peptides would be potential candidates for transient translational pausing under each condition.

## Discussion

We developed a method for the direct isolation and analysis of pep-tRNAs in cells. Pep-tRNAs are “in-between” molecules with both RNA and protein properties. The method developed here exploited the “in-between” property to enrich pep-tRNAs from cellular RNAs (mRNAs, tRNA, and ribosomal RNAs) and proteins that have already completed translation. The organic solvent extraction and silica column purification with high concentrations of guanidium salt were effective for isolating pep-tRNAs. Considering several results, including the RNase treatment, the meta-analysis showing N-terminally-biased peptides derived from pep-tRNAs, and the Tris-conjugated C-terminal ends, we conclude that the method, termed PETEOS, indeed enriches and identifies cellular pep-tRNAs.

A unique feature of the PETEOS method is the direct handling and analysis of pep-tRNAs, providing a complementary approach to other established translatome analyses. Additionally, the PETEOS method has several technical advantages. The time required to halt the cellular processes before the pep-tRNA enrichment is very rapid, since an organic reagent is simply added to the cells. Such quick halting enables the method to capture snapshots of the translation status in the cell, as exemplified by the analyses on heat stress or antibiotic treatments. In addition, the PETEOS procedure is relatively inexpensive and straightforward as compared to ribosome profiling.

The PETEOS method has several limitations. First, the inherently low abundance of pep-tRNAs in the cell makes it difficult to analyze them using LC-MS/MS. Second, the acidic phenol/chloroform extraction, which concentrates proteins and pep-tRNAs while removing RNAs, is not sufficient. Some of the pep-tRNAs move into the aqueous phase, as partly revealed by our systematic separation analysis using model genes (Fig. 1B). Such incomplete separation would lead to unidentified peptidyl-tRNAs, causing some biases in the population and a lack of quantitative assessment. Third, since the peptide moieties derived from pep-tRNAs are mainly identified by tryptic fragments, PETEOS does not have a codon-level resolution, precluding a direct comparison of the method with ribosome profiling. Finally, identifying pep-tRNA-derived peptides, including the C-terminal end of the growing polypeptide, is inherently difficult. The possible reasons for this are as follows. (i) Unless the translation is paused, the C-terminus of pep-tRNA moves continuously, making it difficult to accumulate detectable amounts of the same peptides for MS/MS. (ii) The peptide fragments closer to the N-terminus are, in principle, more abundant (Fig. 2E). These drawbacks are inherent but could be overcome by improving the enrichment efficiency of C-terminal peptides, which are generally considered problematic even in conventional LC-MS/MS-based proteomics.

Despite its incompleteness, PETEOS has potential in that it is complementary to established translatome methods and enables analyses that cannot be achieved by other translatome methods. In addition, although *E. coli* lysates were used as a proof-of-concept experiment, the pep-tRNA analysis by PETEOS could be applicable to any organism.

## Materials and Methods

### *E. coli* strains and plasmids

The *E. coli* BW25113 {genotype: *Δ(araD-araB)567, ΔlacZ4787*(::*rrnB*-3), λ^−^, *rph-1, Δ(rhaD-rhaB)568, hsdR514*} was used as the standard strain. Phage P1-mediated transduction was used to introduce the Δ*tolC* mutation from Keio collection JW5503 (40). The plasmids used in this study are the pCA24N vector and pCA24N-*secM* (from the ASKA library) (41).

### Bacterial growth

*E. coli* cells cultured in LB medium overnight were inoculated in fresh LB medium, and grown at 37 °C with shaking at 180 rpm, unless otherwise noted. The cells were incubated with the antibiotics for 10 min at 37 °C if indicated; final 100 µg/ml of chloramphenicol, 10 µg/mL of puromycin and 50 µg/mL of kasugamycin, or 100 µg/ml of erythromycin, respectively. In case of heat stress experiment, *E. coli* cells were inoculated in LB at 30 °C with shaking at 180 rpm until OD_660_ reached ∼0.4. The culture was then moved into water bath set at 45 °C and incubated for another 30 min with shaking.

*E. coli* cells harboring pCA24N or its derivatives were grown overnight in LB medium containing 20 µg/ml of chloramphenicol, and were then inoculated into fresh LB medium containing chloramphenicol. They were grown at 37 °C with shaking at 180 rpm until the OD_660_ reached ∼0.5. The expression of SecM was then induced by adding 100 μM isopropyl β-D-1-thiogalactopyranoside (IPTG) for 30 min.

### SDS-PAGE under neutral pH conditions and silver staining

Samples were diluted in 2× SDS sample buffer (final 62.5 mM Tris-HCl, pH 6.8, 2% SDS, 10% glycerol, 5% β-mercaptoethanol, a trace of bromophenol blue), as described by Laemmli. SDS sample buffer was treated with RNA Secure (Ambion) according to the manufacturer’s specifications. The samples were separated by the 11% WIDE Range SDS-PAGE system (Nacalai Tesque), as described previously (10, 34).

Silver staining of the gels was performed using a PlusOne Silver Staining Kit, Protein (GE Healthcare), according to the manufacturer’s specifications in protocol “E,” except that glutardialdehyde (25% w/v) was omitted from the *sensitizing solution* in the kit.

### Phenol extraction and TCA/acetone precipitation to fractionate polypeptides and polypeptidyl-tRNAs from nucleic acids

*E. coli* cells grown in 200 mL of LB medium until the mid-log phase were mixed with an approximately equal volume of crushed ice, and collected by centrifugation at 4 °C for 10 min. After supernatant removal, the cell pellet was suspended in 20 mL of TriPURE isolation reagent (Roche) and incubated for 5 min at room temperature. The lysate was then mixed with 4 ml of chloroform, vortexed for 20 sec, incubated for 5 min at room temperature, and centrifuged at 12,000 ×g for 5 min at 4 °C. The colorless to pale yellow transparent upper layer (aqueous phase, 10-12 mL) was removed, leaving an organic phenol phase and a solid middle layer. After the addition of an equal volume (12-15 mL from a 200 mL culture) of ice-cooled 20% TCA/acetone containing 5 mM dithiothreitol (DTT), the mixture was vortexed for 20 sec and incubated for at least 3 hrs at −20 °C. The sample was centrifuged at 15,600 ×g for 20 min at 4 °C, and the supernatant was carefully discarded. The precipitate was vortexed with 10 mL of acetone and centrifuged at 15,600 ×g for 5 min at 4 °C, and then the supernatant was discarded by aspiration. This washing procedure was repeated twice.

### *In vitro* translation

Coupled transcription-translation reaction was performed using PURE*frex 1*.*0* (GeneFrontier) in the presence of 0.2 µM of anti-SsrA oligonucleotide (TTAAGCTGCTAAAGCGTAGTTTTCGTCGTTTGCGACTA) and pre-charged Cy5-Met-tRNA^fMet^ at 37 °C for 40 min (34). DNA templates were prepared by PCR as summarized in Table S2. The reaction was stopped by dilution into twenty volumes of TriPURE isolation reagent. Following phenol-extraction and the precipitation of pep-tRNAs in the aqueous phase were performed according to the manufacturer’s instructions. The pep-tRNAs in the organic phase were precipitated by TCA/acetone precipitation as described above. The precipitates were resolved in 1× SDS sample buffer (62.5 mM Tris-HCl pH 6.8, 2 % SDS, 10% glycerol, 50 mM DTT, treated with RNasecure) and separated by the 11% WIDE Range SDS-PAGE. Then Cy5-labeled pep-tRNAs were detected by using Amersham Typhoon scanner RGB. The signal intensities were quantified by MultiGauge (FUJIFILM) and the ratio of pep-tRNA in the organic phase was calculated as below.

Pep-tRNA (organic phase) / [Pep-tRNA (organic phase) + Pep-tRNA (aqueous phase)]

### Peptidyl-tRNA purification using a silica column

The precipitate obtained from the organic phenol phase was resolved in a 1 mL volume of phase-transfer surfactants (PTS) buffer {100 mM Tris-HCl, pH 6.8, 24 mM sodium deoxycholate, 24 mM sodium N-lauroyl sarcosinate (32)} by vortex mixing.

As the next step, the peptidyl-tRNAs in the PTS buffer were isolated by using a High Pure miRNA Isolation Kit (Roche). After adding 2080 µL of *binding buffer*, the lysate was vortexed and mixed at 37 °C for 30 min with shaking. Then 660 µL of *binding enhancer* was added and the lysate was centrifuged at 12,000 × g for 3 min at 4 °C to remove the precipitates. The supernatant was loaded into 4 columns, and the following steps were performed according to the manufacturer’s instructions. The silica-bound RNA was eluted with 100 µL of PTS buffer, and the collected eluate was divided into two equal portions, and one was treated with 100 µg/mL of RNase A (Promega) at 37 °C for 1 hr. This purification was repeated twice.

### LC-MS/MS sample preparation for the PETEOS experiment

The purified RNA sample in the previous section (100 µL) was mixed with 10 µL of 2 M Tris base (pH= ∼11) and incubated at 80 °C for 20 min to hydrolyze the ester bond of the peptidyl-tRNA. After the addition of 90 µL deionized water, the sample was reduced by the addition of 10 mM DTT at room temperature for 30 min, and then alkylated with 50 mM iodoacetamide in the dark at room temperature for 20 min. The mixture was diluted 5-fold with 50 mM ammonium bicarbonate. For the limited digestion of denatured polypeptides into peptide fragments, 0.5 µg of Lys-C / Trypsin mix (Promega) was added, and the solution was incubated at room temperature for at least 3 hrs. Subsequently, 0.5 µg of Lys-C / Trypsin mix was further added and incubated at 37 °C overnight, whereas for the SecM analysis, 0.5 µg of Lys-C was used. After the digestion, an equal volume of ethyl acetate and 0.5% trifluoroacetic acid (final concentration) was added. The mixture was shaken vigorously for 2 min and centrifuged at 15,700 ×g for 2 min. The upper ethyl acetate layer was removed, and the solvent was removed by a centrifugal evaporator. The residual pellet was dissolved in 0.1% TFA and 2% acetonitrile and desalted, as follows. The solution was applied to an SDB-XC StageTip made by an Empore disk (3M, U.S.A.), equilibrated with 0.1% TFA and 2% acetonitrile, washed with 0.1% TFA and 2% acetonitrile, and eluted with 0.1% TFA and 80 % acetonitrile. The solvent was removed by a centrifugal evaporator, and the residue was dissolved in 0.1% TFA and 2% acetonitrile. This solution was centrifuged at 20,000 ×g for 5 min, and the supernatant was collected and subjected to the LC-MS/MS measurement.

### LC-MS/MS measurement for the PETEOS experiment

The LC-MS/MS measurements were conducted with a Q-Exactive tandem mass spectrometer and an Easy-nLC 1000 nano HPLC system (Thermo Fisher Scientific). The trap column used for the nano HPLC was a 2 cm × 75 μm capillary column packed with 3 μm C18-silica particles (Thermo Fisher Scientific), and the separation column was a 12.5 cm × 75 μm capillary column packed with 3 μm C18-silica particles (Nikkyo Technos, Japan). The flow rate for the nano HPLC was 300 nL/min. The separation was conducted using a 10−40% linear acetonitrile gradient over 70 min, in the presence of 0.1% formic acid. The nanoLC-MS/MS data were acquired in the data-dependent acquisition (DDA) mode controlled by the Xcalibur 4.0 program (Thermo Fisher Scientific). The DDA settings were as follows: the resolution was 70,000 for a full MS scan and 17,500 for an MS2 scan; the AGC target was 3.0E6 for a full MS scan and 5.0E5 for an MS2 scan; the maximum IT was 60 ms for both the full MS and MS2 scans; the full MS scan range was 310-1,500 m/z; and the top 10 signals in each full MS scan were selected for the MS2 scan. The DDA measurement was performed three times for each sample, and two biological replicates were measured in each condition except for the set of antibiotics experiment.

### Data analysis for the PETEOS experiment

The data analysis of the DDA measurements was performed with the Proteome Discoverer 2.4 program bundled with the SEQUEST HT search engine (Thermo Fisher Scientific). The dataset of the amino acid sequences of *E. coli* genes was obtained from the UniProt database (UP000000625, downloaded on Aug. 20^th^, 2019). The ratio calculation and the hypothesis test were conducted by “Summed Abundance Based” and “Individual proteins” mode, respectively. The detailed settings of Proteome Discoverer 2.4 were listed in Supplementary Table S3. Only the peptides/proteins with a < 1 % false discovery rate (by the Percolator algorithm) were selected. For the peptide analyses, the peptides whose number of peptide-spectrum matches (PSMs) was less than three were excluded from the analyses. For the analysis at the protein level, the proteins whose number of detected peptides was less than two were excluded from the analyses. The processes and statistical analyses of the output data of Proteome Discoverer 2.4 were conducted by the R software (version 4.1.3 with in-house scripts).

### Sample preparation and LC-MS/MS measurement and analysis for conventional proteome analysis

The sample preparation was performed according to reference (42). Briefly, the cells were disrupted and containing proteins were precipitated by adding 10% TCA. The proteins were re-dissolved in PTS buffer and alkylated with 55 mM iodoacetamide after reduction with 10 mM DTT. Then, the proteins were digested by Trypsin/Lys-C Mix (Promega). After the digestion, surfactants were removed by the ethyl acetate treatment at low pH, and the resulting peptides were desalted with an SDB-XC StageTip.

The measurement for the comparison against the phenol-extracted proteins (Figure S1A) was performed in DDA mode using a Q-Exactive tandem mass spectrometer and an Easy-nLC 1000 nano HPLC system (Thermo Fisher Scientific). The detailed settings were as described above. The data analysis was performed using Proteome Discoverer 2.4 with the same settings as described above. For the heat shock experiment, the measurement was performed in DIA/SWATH-acquisition mode using a TripleTOF 6600 tandem mass spectrometer and an Eksigent nanoLC 415 nano HPLC system (SCIEX). The analysis of DIA/SWATH data was performed using DIA-NN software (version 1.7.16) with default settings (43). The library for DIA-SWATH analysis was obtained from the SWATHatlas database (http://www.swathatlas.org/, accessed on 30 April 2021); the original data were acquired by Midha *et al* (44). The following analysis was performed using in-house R scripts (R.app for Mac, version 4.1.3). The detailed settings of the measurement and the analyses were the same as described in the previous report (42).

## Data Availability Statement

Data in this manuscript have been uploaded to the Mendeley Dataset public repository (https://doi.org/10.17632/xzycy8m6tb.1). The mass spectrometry data have been deposited in the jPOST repository (45) (https://repository.jpostdb.org/), with reference number JPST001572/PXD033726.

## Author contributions

YC, AY and AK performed experiments; TN and AY performed LC-MS/MS and analyzed the MS data; YC, TN, and HT conceived the study, designed experiments, and analyzed the results; YC, TN, and HT supervised the entire project; TN, AY, YC and HT wrote the manuscript.

## Acknowledgments

We thank Eri Uemura for technical support; the Bio-support Center at Tokyo Tech for DNA sequencing; and the Cell Biology Center Research Core Facility at Tokyo Tech for the Q-Exactive and TripleTOF 6600 mass spectrometry measurements. This work was supported by MEXT Grants-in-Aid for Scientific Research (Grant Numbers 17K15073 to TN, 17K15062, 19K16038 to YC, JP26116002, JP18H03984, JP20H05925 to HT) and a grant from the Ohsumi Frontier Science Foundation to YC.

## References

1. Vogel, E. M. Marcotte, Insights into the regulation of protein abundance from proteomic and transcriptomic analyses. Nature Reviews Genetics 13, 227–232 (2012).

2. S. Pechmann, F. Willmund, J. Frydman, The Ribosome as a Hub for Protein Quality Control. Molecular Cell 49, 411–421 (2013).

3. K. Ito, S. Chiba, Arrest peptides: Cis-acting modulators of translation. Annual Review of Biochemistry 82, 171–202 (2013).

4. K. C. Stein, J. Frydman, The stop-and-go traffic regulating protein biogenesis: How translation kinetics controls proteostasis. Journal of Biological Chemistry 294, 2076–2084 (2019).

5. E. Samatova, J. Daberger, M. Liutkute, M. v Rodnina, Translational Control by Ribosome Pausing in Bacteria: How a Non-uniform Pace of Translation Affects Protein Production and Folding. Frontiers in Microbiology 11, 619430 (2021).

6. D. N. Wilson, S. Arenz, R. Beckmann, Translation regulation via nascent polypeptide-mediated ribosome stalling. Current Opinion in Structural Biology 37, 123–133 (2016).

7. J. Choi, et al., How Messenger RNA and Nascent Chain Sequences Regulate Translation Elongation. Annu Rev Biochem 87, 421–449 (2018).

8. H. Nakatogawa, K. Ito, Secretion monitor, secM, undergoes self-translation arrest in the cytosol. Molecular Cell 7, 185–192 (2001).

9. H. Nakatogawa, K. Ito, The ribosomal exit tunnel functions as a discriminating gate. Cell 108, 629–36 (2002).

10. Y. Chadani, T. Niwa, S. Chiba, H. Taguchi, K. Ito, Integrated in vivo and in vitro nascent chain profiling reveals widespread translational pausing. Proc Natl Acad Sci U S A 113, E829–E838 (2016).

11. S. Iwasaki, N. T. Ingolia, The Growing Toolbox for Protein Synthesis Studies. Trends Biochem Sci 42, 612–624 (2017).

12. N. T. Ingolia, S. Ghaemmaghami, J. R. S. Newman, J. S. Weissman, Genome-wide analysis in vivo of translation with nucleotide resolution using ribosome profiling. Science 324, 218–223 (2009).

13. N. T. Ingolia, G. A. Brar, S. Rouskin, A. M. McGeachy, J. S. Weissman, The ribosome profiling strategy for monitoring translation in vivo by deep sequencing of ribosome-protected mRNA fragments. Nature Protocols 7, 1534–1550 (2012).

14. G. A. Brar, J. S. Weissman, Ribosome profiling reveals the what, when, where and how of protein synthesis. Nature Reviews Molecular Cell Biology 16, 651–664 (2015).

15. N. T. Ingolia, J. A. Hussmann, J. S. Weissman, Ribosome profiling: Global views of translation. Cold Spring Harbor Perspectives in Biology 11, a032698 (2019).

16. A. R. Buskirk, R. Green, Ribosome pausing, arrest and rescue in bacteria and eukaryotes. Philosophical Transactions of the Royal Society B: Biological Sciences 372, 20160183 (2017).

17. F. Mohammad, C. J. Woolstenhulme, R. Green, A. R. Buskirk, Clarifying the Translational Pausing Landscape in Bacteria by Ribosome Profiling. Cell Reports 14, 686–694 (2016).

18. D. C. Dieterich, A. J. Link, J. Graumann, D. A. Tirrell, E. M. Schuman, Selective identification of newly synthesized proteins in mammalian cells using bioorthogonal noncanonical amino acid tagging (BONCAT). Proc Natl Acad Sci U S A 103, 9482–9487 (2006).

19. A. J. M. Howden, et al., QuaNCAT: Quantitating proteome dynamics in primary cells. Nature Methods 10, 343–346 (2013).

20. R. Aviner, T. Geiger, O. Elroy-Stein, Genome-wide identification and quantification of protein synthesis in cultured cells and whole tissues by puromycin-associated nascent chain proteomics (PUNCH-P). Nature Protocols 9, 751–760 (2014).

21. R. Aviner, T. Geiger, O. Elroy-Stein, Novel proteomic approach (PUNCH-P) reveals cell cycle-specific fluctuations in mRNA translation. Genes and Development 27, 1834–1844 (2013).

22. C. M. Forester, et al., Revealing nascent proteomics in signaling pathways and cell differentiation. Proc Natl Acad Sci U S A 115, 2353–2358 (2018).

23. J. Uchiyama, et al., pSNAP: Proteome-wide analysis of elongating nascent polypeptide chains. iScience 25, 104516 (2022).

24. H. Muto, H. Nakatogawa, K. Ito, Genetically Encoded but Nonpolypeptide Prolyl-tRNA Functions in the A Site for SecM-Mediated Ribosomal Stall. Molecular Cell 22, 545–552 (2006).

25. F. Gong, K. Ito, Y. Nakamura, C. Yanofsky, The mechanism of tryptophan induction of tryptophanase operon expression: Tryptophan inhibits release factor-mediated cleavage of TnaC-peptidyl-tRNAPro. Proc Natl Acad Sci U S A 98, 8997–9001 (2001).

26. H. Onouchi, et al., Nascent peptide-mediated translation elongation arrest coupled with mRNA degradation in the CGS1 gene of Arabidopsis. Genes and Development 19, 1799–1810 (2005).

27. M. Dermit, M. Dodel, F. K. Mardakheh, Methods for monitoring and measurement of protein translation in time and space. Molecular BioSystems 13, 2477–2488 (2017).

28. R. Milo, P. Jorgensen, U. Moran, G. Weber, M. Springer, BioNumbers The database of key numbers in molecular and cell biology. Nucleic Acids Research 38, D750–3 (2010).

29. K. Ito, et al., Nascentome analysis uncovers futile protein synthesis in Escherichia coli. PLoS ONE 6, e28413 (2011).

30. R. Boom, et al., Rapid and simple method for purification of nucleic acids. Journal of Clinical Microbiology 28, 495–503 (1990).

31. B. Vogelstein, D. Gillespie, Preparative and analytical purification of DNA from agarose. Proc Natl Acad Sci U S A 76, 615–619 (1979).

32. T. Masuda, M. Tomita, Y. Ishihama, Phase transfer surfactant-aided trypsin digestion for membrane proteome analysis. Journal of Proteome Research 7, 731–740 (2008).

33. Y. Shimizu, et al., Cell-free translation reconstituted with purified components. Nature Biotechnology 19, 751–755 (2001).

34. Y. Chadani, et al., Intrinsic Ribosome Destabilization Underlies Translation and Provides an Organism with a Strategy of Environmental Sensing. Molecular Cell 68, 528–539.e5 (2017).

35. C. P. Flessel, P. Ralph, A. Rich, Polyribosomes of Growing Bacteria. Science 158, 658–660 (1967).

36. D. W. Allen, P. C. Zamecnik, The effect of puromycin on rabbit reticulocyte ribosomes. Biochim Biophys Acta 55, 865–874 (1962).

37. K. Kannan, et al., The general mode of translation inhibition by macrolide antibiotics. Proc Natl Acad Sci U S A 111, 15958–15963 (2014).

38. A. R. Davis, D. W. Gohara, M. N. F. Yap, Sequence selectivity of macrolide-induced translational attenuation. Proc Natl Acad Sci U S A 111, 15379–15384 (2014).

39. K. Saito, et al., Ribosome collisions induce mRNA cleavage and ribosome rescue in bacteria. Nature 603, 503–508 (2022).

40. T. Baba, et al., Construction of Escherichia coli K-12 in-frame, single-gene knockout mutants: The Keio collection. Molecular Systems Biology 2, 0008 (2006).

41. M. Kitagawa, et al., Complete set of ORF clones of Escherichia coli ASKA library (A complete set of E. coli K-12 ORF archive): unique resources for biological research. DNA Research 12, 291–299 (2005).

42. H. Onodera, T. Niwa, H. Taguchi, Y. Chadani, Novel trans-translation-associated gene regulation revealed by prophage excision-triggered switching of ribosome rescue pathway. bioRxiv, 2022.04.27.489667 (2022).

43. V. Demichev, C. B. Messner, S. I. Vernardis, K. S. Lilley, M. Ralser, DIA-NN: neural networks and interference correction enable deep proteome coverage in high throughput. Nat Methods 17, 41–44 (2020).

44. M. K. Midha, et al., A comprehensive spectral assay library to quantify the Escherichia coli proteome by DIA/SWATH-MS. Sci Data 7 (2020).

45. S. Okuda, et al., jPOSTrepo: an international standard data repository for proteomes. Nucleic Acids Res 45, D1107–D1111 (2017).

