## Supplemental Figures and Tables for "A method to enrich polypeptidyl-tRNAs to capture snapshots of translation in the cell"

**Supplementary Figure S1**: Numbers of MS-identified proteins prepared by a conventional method and an organic extraction.

**Supplementary Figure S2**: Properties of the peptides/proteins identified by the PETEOS method.

**Supplementary Figure S3**: Investigation of the changes in the nascent peptides/proteins by heat-shock and antibiotics.

**Supplementary Table S1**: The detected peptides whose C-terminus are not lysine nor arginine under each condition.

**Supplementary Table S2**: Preparation of the templated DNA fragments used for in vitro translation.

**Supplementary Table S3**: Settings of Proteome Discoverer 2.4.

**Supplementary Dataset S1**: List of identified peptides and proteins by the PETEOS method in replicate 1 and 2 (10 sheets).

**Supplementary Dataset S2**: List of identified peptides and proteins by the PETEOS method in the heat-shock experiment (two replicates, 14 sheets).

**Supplementary Dataset S3**: Total proteome changes by the heat-shock evaluated by the SWATH-MS acquisition method (two replicates, one sheet).

**Supplementary Dataset S4**: List of identified peptides and proteins by the PETEOS method in the antibiotic-treatment experiment (17 sheets).

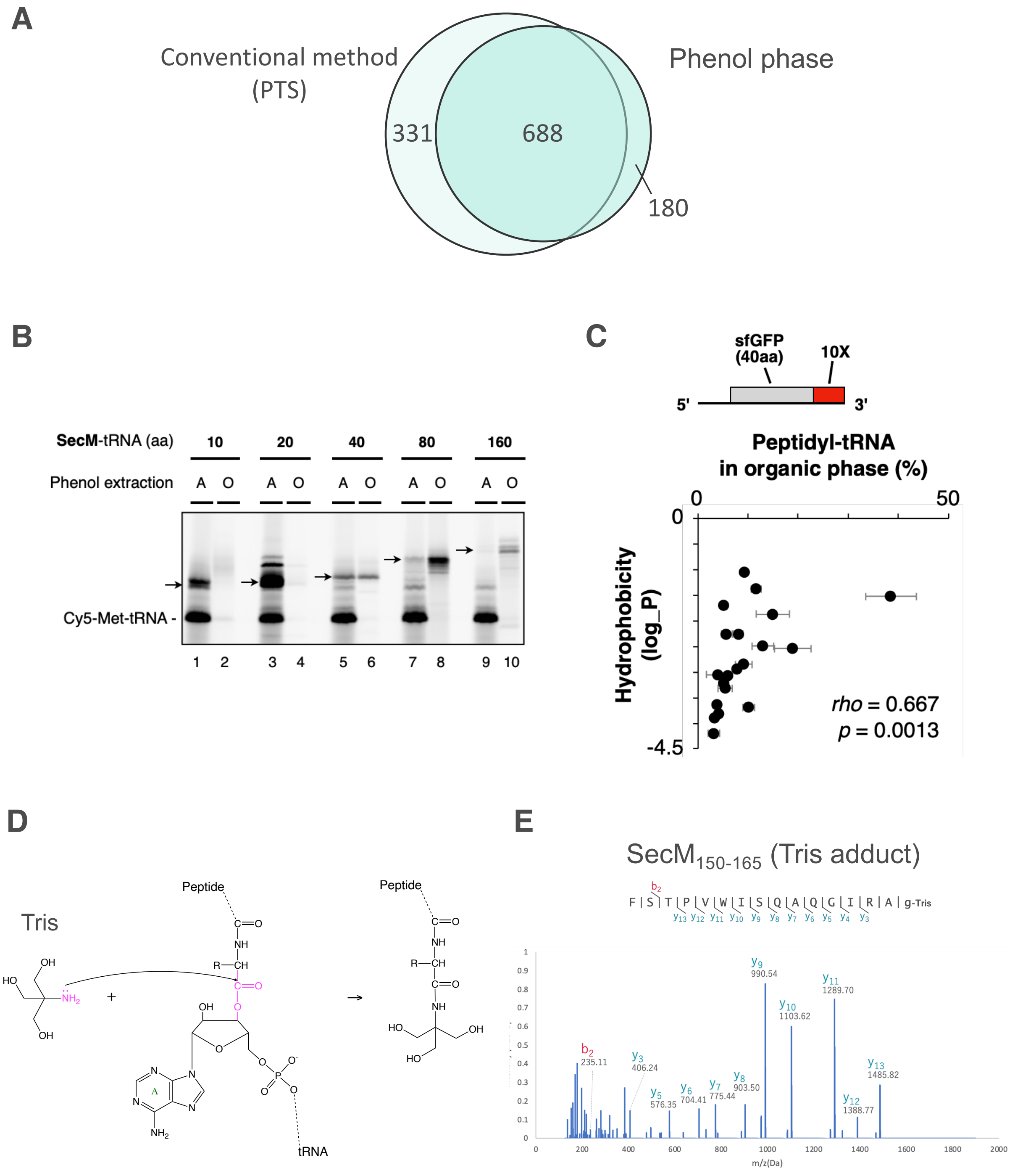


**Figure S1. Numbers of MS-identified proteins prepared by a conventional method and an organic extraction.**

**A.** For the conventional method, phase-transfer surfactants (PTS) lysis buffer was used to extract the proteins from the cell. For the organic extraction, the proteins in the phenol/chloroform phase, Step 1 of the PETEOS workflow, were identified by LC-MS/MS. The numbers of proteins identified by LC-MS/MS with two or more peptides are shown.

**B**. Representative gel image of the fractionation of SecM-tRNAs by phenol-extraction. In vitro translation mixtures of the various length of *secM* nonstop mRNA were phenol-extracted. The pep-tRNA in the aqueous (A, odd-numbered lanes) or organic (O, even-numbered lanes) phase were analyzed in parallel. The signals used for the quantification were indicated by arrows.

**C**. Contribution of the hydrophobicity in phenol-based fractionation. The nonstop mRNAs carrying N-terminal 40 a.a. part of sfGFP and 20 kinds of 10 consecutive a.a. sequence (10X) were individually translated by PURE*frex* and analyzed as shown in Fig. 1B. The quantified values and the hydrophobicity (logP) of the 10X sequence were represented by two-dimensional plot. Spearman's rho and P-values calculated by Spearman's rank correlation tests are also shown.

**D**. The nucleophilic substitution reaction replaces the nucleic acid moiety with tris(hydroxymethyl)aminomethane (Tris)-base.

**E**. MS2 spectra of the C-terminal peptides of SecM modified by Tris-base ($_{150}$FSTPVWISQAQGIRAG$_{165}$-Tris) from the *E. coli* cells overexpressed SecM.

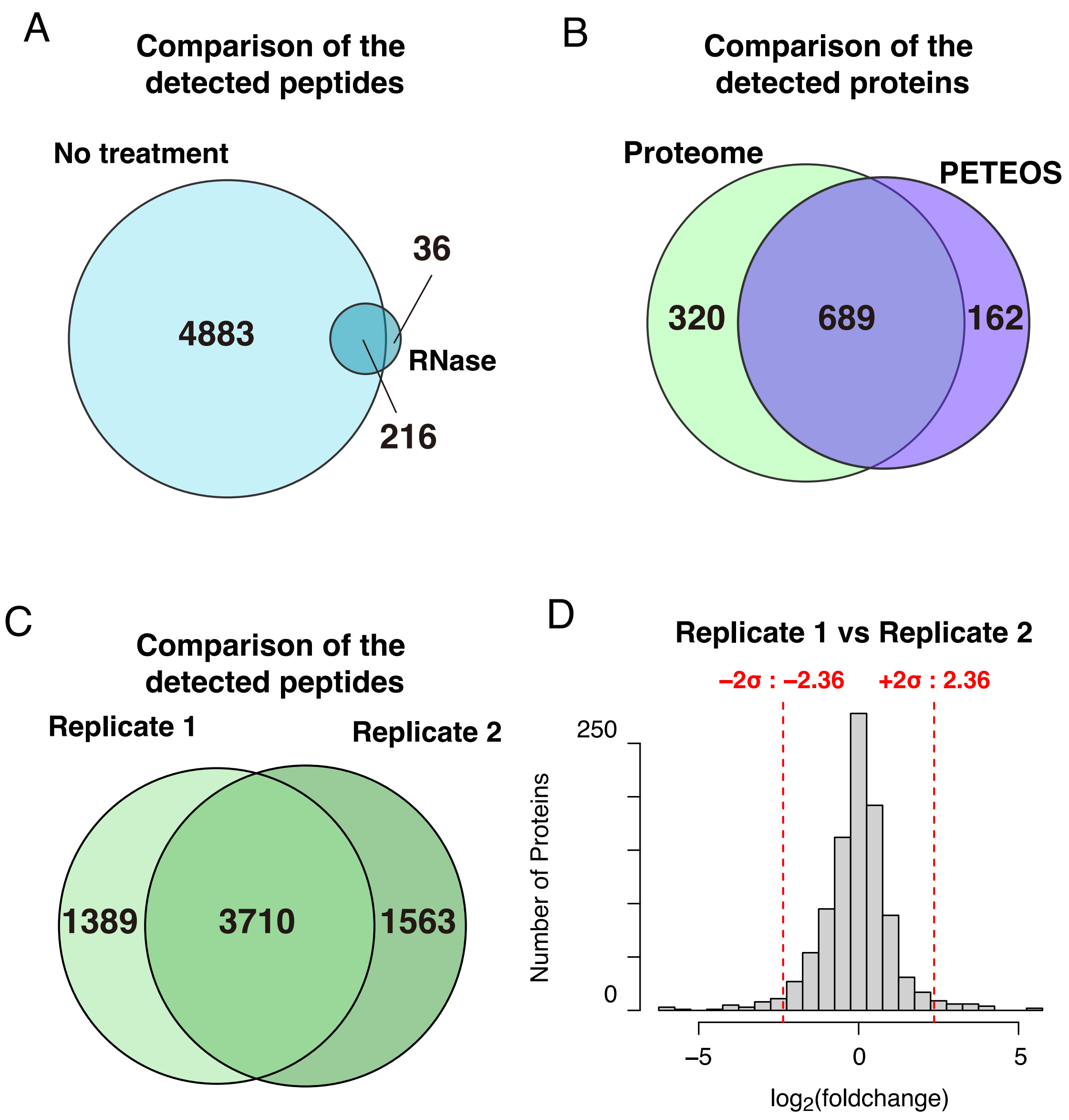


**Figure S2. Properties of the peptides/proteins identified by the PETEOS method.**

**A.** Number of the detected peptides by the PETEOS method. As a control, the number of peptides detected in the RNase-treated fraction and its overlap were also shown. The peptides with less than three number of PSMs were omitted.

**B.** Overlap of the proteins detected by the PETEOS method and a conventional proteome method. The proteins with < 1% FDR and ≥ 2 number of detected peptides were counted.

**C.** Number of peptides obtained from replicates 1 and 2 and their overlap. The peptides with less than three number of PSMs were omitted.

**D.** Quantitative assessment of the proteins between replicate 1 and 2. The histogram showed the distribution of the fold change values between replicate 1 and 2. The fold change values were calculated by Proteome Discoverer 2.4 (detailed settings were described in the Materials and Methods section and Supplementary Table S3). The fold change values were normalized by the median value. Red dotted lines represent ± 2$\sigma$ points of the distribution (± 2.36 as $\log_2$ value, indicating ~5.1 or ~1/5.1 fold changes).

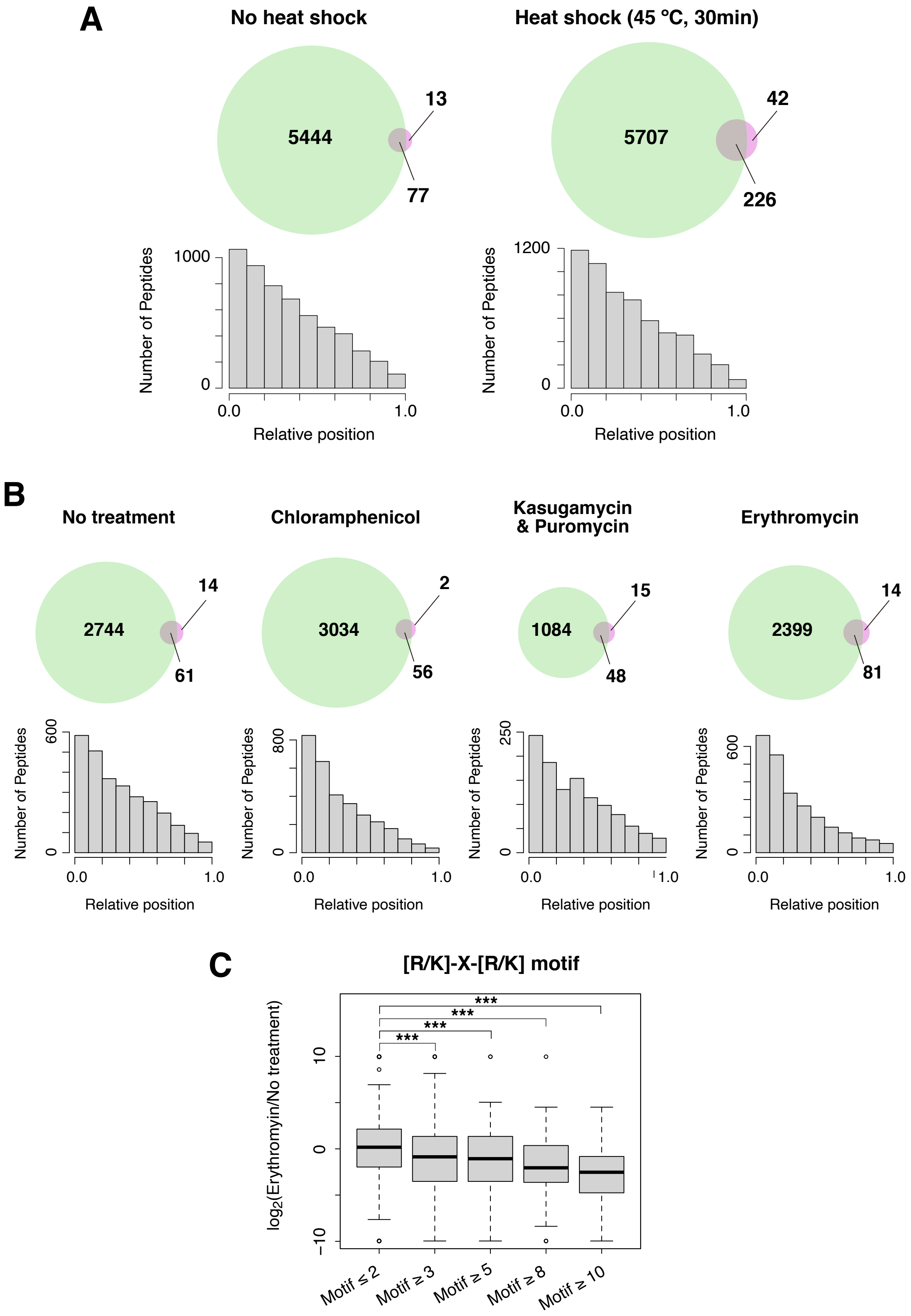


**Figure S3. Investigation of the changes in the nascent peptides/proteins by heat-**

**shock and antibiotics.**

**A.** (Upper) Number of the detected peptides from the samples for Heat shock comparison. A green circle indicates the detected peptides from the no RNase-treated fraction, and magenta indicates those from the RNase-treated fraction. The peptides with less than three number of PSMs were omitted. (Lower) Meta-analysis of the relative position of the peptides identified in each fraction.

**B.** (Upper) Number of the detected peptides from the samples treated with antibiotics. Green circle indicates the detected peptides from the no RNase-treaed fraction and magenta indicates those from the RNase-treated fraction. The peptides with less than three number of PSMs were omitted. (Lower) Meta-analysis of the relative position of the peptides identified in each fraction.

**C.** Distribution of the fold changes by the treatment of Erythromycin. The box portions and the central bands are described according to the 25th percentile and the median, respectively. The number of proteins in each group was as follows; $n$ =589 for Motif ≤ 2, $n$ =486 for Motif ≥3, $n$ =226 for Motif ≥5, $n$ =78 for Motif ≥8, and $n$ = 33 for Motif ≥10. *** $P < 0.0005$, by Welch's t-test.

Supplementary Table S1 The detected peptides whose C-terminus is not lysine nor arginine under each condition.

| UniProt ID | No treatment | | | | | Chloramphenicol | | | | | Kasugamycin & Puromycin | | | | | Erythromycin | | | | |
|---|---|---|---|---|---|---|---|---|---|---|---|---|---|---|---|---|---|---|---|---|
| | All_seq | Number of Detected Peptides | Number of Detected Tris-added peptides | Detected Position | Sum of corre-sponding PSM numbers | All_seq | Number of Detected Peptides | Number of Detected Tris-added peptides | Detected Position | Sum of corre-sponding PSM numbers | All_seq | Number of Detected Peptides | Number of Detected Tris-added peptides | Detected Position | Sum of corre-sponding PSM numbers | All_seq | Number of Detected Peptides | Number of Detected Tris-added peptides | Detected Position | Sum of corre-sponding PSM numbers |
| P00579 | | | | | | | | | | | | | | | | LSDLITGFVD<br>LSDLITGFVDPNAEEDLAPT<br>LSDLITGFVDPNAEEDLAPTATHVGSELSQEDLDDDEDEDEED<br>LSDLITGFVDPNAEEDLAPTATHVGSELSQEDLDDDEDEDEEDGD<br>\|LSDLITGFVDPNAEEDLAPTATHVGSELSQEDLDDDEDEDEEDGDD | 5 | 0 | 158-167<br>158-177<br>158-200<br>158-202<br>158-203 | 10 |
| P00805 | | | | | | | | | | | | | | | | VGVENLVNAVPQL<br>VGVENLVNAVPQLKD | 2 | 1 | 52-64<br>52-66 | 12 |
| P02358 | | | | | | | | | | | | | | | | AHYVLMNVEAPQEVID<br>AHYVLMNVEAPQEVIDE<br>AHYVLMNVEAPQEVIDEL<br>AHYVLMNVEAPQEVIDELE | 4 | 3 | 57-72<br>57-73<br>57-74<br>57-75 | 31 |
| P06959 | | | | | | KEAAPAAAP<br>KEAAPAAAPAAAAA | 2 | 1 | 92-100<br>92-105 | 9 | | | | | | | | | | |
| P06996 | VLSLLVPALLVA<br>VLSLLVPALLVAGA<br>VLSLLVPALLVAGAA<br>VLSLLVPALLVAGAAN<br>VLSLLVPALLVAGAANA<br>VLSLLVPALLVAGAANAAE<br>VLSLLVPALLVAGAANAAEVY<br>VLSLLVPALLVAGAANAAEVYNKD<br>VLSLLVPALLVAGAANAAEVYNKDG<br>VLSLLVPALLVAGAANAAEVYNKDGN | 10 | 1 | 5-16<br>5-18<br>5-19<br>5-20<br>5-21<br>5-23<br>5-25<br>5-28<br>5-29<br>5-30 | 84 | VLSLLVPALL<br>VLSLLVPALLVA<br>VLSLLVPALLVAG<br>VLSLLVPALLVAGA<br>VLSLLVPALLVAGAA<br>VLSLLVPALLVAGAAN<br>VLSLLVPALLVAGAANA<br>VLSLLVPALLVAGAANAA<br>VLSLLVPALLVAGAANAAE<br>VLSLLVPALLVAGAANAAEVY<br>VLSLLVPALLVAGAANAAEVYN<br>VLSLLVPALLVAGAANAAEVYNKD<br>VLSLLVPALLVAGAANAAEVYNKDG<br>VLSLLVPALLVAGAANAAEVYNKDGN | 14 | 4 | 5-14<br>5-16<br>5-17<br>5-18<br>5-19<br>5-20<br>5-21<br>5-22<br>5-23<br>5-25<br>5-26<br>5-28<br>5-29<br>5-30 | 239 | VLSLLVPALLVAGAAN<br>VLSLLVPALLVAGAANAAEVY<br>VLSLLVPALLVAGAANAAEVYNKD<br>VLSLLVPALLVAGAANAAEVYNKDG | 4 | 1 | 5-20<br>5-25<br>5-28<br>5-29 | 41 | VLSLLVPALLVAGAAN<br>VLSLLVPALLVAGAANAAEVY | 2 | 1 | 5-20<br>5-25 | 27 |
| | GETQVTDQLTGY<br>GETQVTDQLTGYGQWEY | 2 | 0 | 63-74<br>63-79 | 5 | GETQVTDQLTGYGQ<br>GETQVTDQLTGYGQWE | 2 | 0 | 63-76<br>63-78 | 6 | | | | | | | | | | |
| | NYGVVYDVTSWTDVLPE<br>NYGVVYDVTSWTDVLPEFGGD<br>NYGVVYDVTSWTDVLPEFGGDTY | 3 | 1 | 114-130<br>114-134<br>114-136 | 10 | NYGVVYDVTSW<br>NYGVVYDVTSWTDVLPE<br>NYGVVYDVTSWTDVLPEFGG<br>NYGVVYDVTSWTDVLPEFGGD<br>NYGVVYDVTSWTDVLPEFGGDTYGS<br>NYGVVYDVTSWTDVLPEFGGDTYGSD | 6 | 3 | 114-124<br>114-130<br>114-133<br>114-134<br>114-138<br>114-139 | 29 | NYGVVYDVTSWTDVLPE<br>NYGVVYDVTSWTDVLPEFGGD<br>NYGVVYDVTSWTDVLPEFGGDTY | 3 | 0 | 114-130<br>114-134<br>114-136 | 9 | | | | | |
| P09169 | | | | | | LLGIVLTTPIAISS<br>LLGIVLTTPIAISSF<br>LLGIVLTTPIAISSFAS | 3 | 0 | 5-18<br>5-19<br>5-21 | 9 | | | | | | | | | | |
| P09373 | | | | | | THAPVDFDTAVAS<br>THAPVDFDTAVASTI<br>THAPVDFDTAVASTITS | 3 | 0 | 68-80<br>68-82<br>68-84 | 9 | | | | | | | | | | |
| P0A6R3 | | | | | | | | | | | | | | | | NYFAQLNGQDVNDLY<br>NYFAQLNGQDVNDLYELVLAE | 2 | 0 | 37-51<br>37-57 | 5 |
| P0A7A9 | LVAVPH<br>LVAVPHS | 2 | 0 | 106-111<br>106-112 | 7 | | | | | | | | | | | | | | | |
| P0A7F3 | | | | | | IENTFLSEDQVDQLAL<br>IENTFLSEDQVDQLALYAP | 2 | 0 | 61-76<br>61-79 | 5 | | | | | | | | | | |
| P0A7K2 | | | | | | | | | | | | | | | | DLVESAPAAL<br>DLVESAPAALKE<br>DLVESAPAALKEG<br>DLVESAPAALKEGV<br>DLVESAPAALKEGVS<br>DLVESAPAALKEGVSKDD | 6 | 0 | 86-95<br>86-97<br>86-98<br>86-99<br>86-100<br>86-103 | 19 |
| P0A853 | SGMNPFLLDSEDVFID<br>SGMNPFLLDSEDVFIDL<br>SGMNPFLLDSEDVFIDLLTDSGTG<br>SGMNPFLLDSEDVFIDLLTDSGTGAVTQS<br>SGMNPFLLDSEDVFIDLLTDSGTGAVTQSMQAA | 5 | 3 | 34-49<br>34-50<br>34-57<br>34-62<br>34-66 | 12 | SGMNPFLLDSE<br>SGMNPFLLDSEDVFIDLLTDSGTGAVTQ<br>SGMNPFLLDSEDVFIDLLTDSGTGAVTQS<br>SGMNPFLLDSEDVFIDLLTDSGTGAVTQSM<br>SGMNPFLLDSEDVFIDLLTDSGTGAVTQSMQA<br>SGMNPFLLDSEDVFIDLLTDSGTGAVTQSMQAA<br>SGMNPFLLDSEDVFIDLLTDSGTGAVTQSMQAAM<br>SGMNPFLLDSEDVFIDLLTDSGTGAVTQSMQAAMMRG | 8 | 1 | 34-44<br>34-61<br>34-62<br>34-63<br>34-65<br>34-66<br>34-67<br>34-70 | 25 | | | | | | | | | | |
| P0A870 | | | | | | | | | | | | | | | | LYQPQDATTNPSLIL<br>LYQPQDATTNPSLILNAAQIP<br>LYQPQDATTNPSLILNAAQIPEY | 3 | 1 | 26-40<br>26-46<br>26-48 | 22 |
| P0A8F4 | | | | | | | | | | | | | | | | EQVGDEHIGVIPEDCY<br>EQVGDEHIGVIPEDCYY<br>EQVGDEHIGVIPEDCYYKD | 3 | 1 | 34-49<br>34-50<br>34-52 | 16 |
| P0A905 | | | | | | SLATAAGAVAGGVAG<br>SLATAAGAVAGGVAGQGVQS | 2 | 1 | 85-99<br>85-104 | 8 | | | | | | SLATAAGAVAGGVAGQ<br>SLATAAGAVAGGVAGQGVQS | 2 | 1 | 85-100<br>85-104 | 8 |

(Supplementary Table S1, continued)

| UniProt ID | No treatment | | | | | Chloramphenicol | | | | | Kasugamycin & Puromycin | | | | | Erythromycin | | | | |
|---|---|---|---|---|---|---|---|---|---|---|---|---|---|---|---|---|---|---|---|---|
| | All_seq | Number of Detected Peptides | Number of Detected Tris-added peptides | Detected Position | Sum of corre-sponding PSM numbers | All_seq | Number of Detected Peptides | Number of Detected Tris-added peptides | Detected Position | Sum of corre-sponding PSM numbers | All_seq | Number of Detected Peptides | Number of Detected Tris-added peptides | Detected Position | Sum of corre-sponding PSM numbers | All_seq | Number of Detected Peptides | Number of Detected Tris-added peptides | Detected Position | Sum of corre-sponding PSM numbers |
| P0A910 | | | | | | LGWSQYHDTGFINNNGPTHENQLGAGAFGG<br>LGWSQYHDTGFINNNGPTHENQLGAGAFGGY | 2 | 0 | 34-63<br>34-64 | 3 | LGWSQYHDTGFINNNGPTHENQLGAGAFGGY<br>LGWSQYHDTGFINNNGPTHENQLGAGAFGGYQVNPY | 2 | 0 | 34-64<br>34-69 | 6 | LGWSQYHDTGFI<br>LGWSQYHDTGFINN<br>LGWSQYHDTGFINNN<br>LGWSQYHDTGFINNNG<br>LGWSQYHDTGFINNNGPTH<br>LGWSQYHDTGFINNNGPTHE<br>LGWSQYHDTGFINNNGPTHEN<br>LGWSQYHDTGFINNNGPTHENQL<br>LGWSQYHDTGFINNNGPTHENQLGAGAF<br>LGWSQYHDTGFINNNGPTHENQLGAGAFG<br>LGWSQYHDTGFINNNGPTHENQLGAGAFGGY<br>LGWSQYHDTGFINNNGPTHENQLGAGAFGGYQVNPY | 12 | 1 | 34-45<br>34-47<br>34-48<br>34-49<br>34-52<br>34-53<br>34-54<br>34-56<br>34-61<br>34-62<br>34-64<br>34-69 | 112 |
| | | | | | | | | | | | | | | | | LGYPITDDLDIY<br>LGYPITDDLDIYTRL | 2 | 1 | 104-115<br>104-118 | 11 |
| | | | | | | | | | | | LEYQWTNNIGDAH<br>LEYQWTNNIGDAHTIGTRPD | 2 | 0 | 160-172<br>160-179 | 6 | LEYQWTNNIGDAH<br>LEYQWTNNIGDAHTIGTRPD | 2 | 1 | 160-172<br>160-179 | 21 |
| | ATLKPEGQAA<br>ATLKPEGQAALDQLYS<br>ATLKPEGQAALDQLYSQLS<br>ATLKPEGQAALDQLYSQLSNLD | 4 | 1 | 228-237<br>228-243<br>228-246<br>228-249 | 10 | ATLKPEGQAALDQLYS<br>ATLKPEGQAALDQLYSQLS<br>ATLKPEGQAALDQLYSQLSNLD | 3 | 1 | 228-243<br>228-246<br>228-249 | 18 | ATLKPEGQAA<br>ATLKPEGQAALDQL<br>\|ATLKPEGQAALDQLYSQLSNLD | 3 | 0 | 228-237<br>228-242<br>228-249 | 11 | ATLKPEGQ<br>ATLKPEGQAALDQLY<br>ATLKPEGQAALDQLYS | 3 | 1 | 228-235<br>228-242<br>228-243 | 31 |
| P0A915 | AGSATVRPTEGAGGTLGSLGGF<br>AGSATVRPTEGAGGTLGSLGGFS | 2 | 0 | 31-52<br>31-53 | 4 | AGSATVRPTEGAGGTLGS<br>AGSATVRPTEGAGGTLGSLGG<br>AGSATVRPTEGAGGTLGSLGGF<br>AGSATVRPTEGAGGTLGSLGGFS<br>AGSATVRPTEGAGGTLGSLGGFSV<br>AGSATVRPTEGAGGTLGSLGGFSVTNNTQLGL | 6 | 3 | 31-48<br>31-51<br>31-52<br>31-53<br>31-54<br>31-62 | 21 | | | | | | AGSATVRP<br>AGSATVRPTE<br>AGSATVRPTEG<br>AGSATVRPTEGAG<br>AGSATVRPTEGAGGTL<br>AGSATVRPTEGAGGTLG<br>AGSATVRPTEGAGGTLGS<br>AGSATVRPTEGAGGTLGSL<br>AGSATVRPTEGAGGTLGSLGG<br>AGSATVRPTEGAGGTLGSLGGF<br>AGSATVRPTEGAGGTLGSLGGFS | 11 | 5 | 31-38<br>31-40<br>31-41<br>31-43<br>31-46<br>31-47<br>31-48<br>31-49<br>31-51<br>31-52<br>31-53 | 38 |
| | | | | | | ATGDIATVHHLPPTLMAQW<br>ATGDIATVHHLPPTLMAQWYFGDASS | 2 | 1 | 90-108<br>90-115 | 5 | | | | | | | | | | |
| P0A917 | | | | | | IACLSALAAVL<br>IACLSALAAVLAF<br>IACLSALAAVLAFTAGTS<br>IACLSALAAVLAFTAGTSVAATS | 4 | 2 | 4-14<br>4-16<br>4-21<br>4-26 | 16 | | | | | | IACLSALAAVL<br>IACLSALAAVLAF<br>IACLSALAAVLAFTAGTSVAATS | 3 | 1 | 4-14<br>4-16<br>4-26 | 14 |
| | | | | | | YRYEEDNSPLGV<br>YRYEEDNSPLGVIGS<br>YRYEEDNSPLGVIGSF<br>YRYEEDNSPLGVIGSFTY<br>YRYEEDNSPLGVIGSFTYTE | 5 | 2 | 51-62<br>51-65<br>51-66<br>51-68<br>51-70 | 18 | | | | | | | | | | |
| P0A953 | | | | | | RAVITGLGIVSSIGNNQQEVLAS<br>RAVITGLGIVSSIGNNQQEVLASL | 2 | 2 | 3-25<br>3-26 | 8 | | | | | | | | | | |
| P0A998 | | | | | | INTVESPFAE<br>INTVESPFAEYSS<br>INTVESPFAEYSSLDELFQETY | 3 | 1 | 70-79<br>70-82<br>70-91 | 17 | | | | | | | | | | |
| P0A9Q5 | | | | | | LHSLLDEGSLVELGS<br>LHSLLDEGSLVELGSE | 2 | 1 | 61-75<br>61-76 | 9 | | | | | | | | | | |
| P0ABI4 | | | | | | | | | | | | | | | | LEVEESQPLVNAVWIDLVEPD<br>LEVEESQPLVNAVWIDLVEPDDDERL | 2 | 1 | 15-35<br>15-40 | 12 |
| P0ABU9 | | | | | | | | | | | | | | | | RDNLTGSEQIFY<br>RDNLTGSEQIFYS | 2 | 1 | 73-84<br>73-85 | 14 |
| P0ACA3 | | | | | | | | | | | | | | | | SVMTLFSGPTDIY<br>SVMTLFSGPTDIYSH | 2 | 0 | 9-21<br>9-23 | 7 |
| P0ACF4 | ALDAIIASVTESL<br>ALDAIIASVTESLKE<br>ALDAIIASVTESLKEG<br>ALDAIIASVTESLKEGDD<br>ALDAIIASVTESLKEGDDV<br>ALDAIIASVTESLKEGDDVA<br>ALDAIIASVTESLKEGDDVAL<br>ALDAIIASVTESLKEGDDVALVG | 8 | 1 | 24-36<br>24-38<br>24-39<br>24-41<br>24-42<br>24-43<br>24-44<br>24-46 | 38 | ALDAIIASVTES<br>ALDAIIASVTESLKEGD<br>ALDAIIASVTESLKEGDDVAL | 3 | 2 | 24-35<br>24-40<br>24-44 | 28 | ALDAIIASVTESLKEGD<br>ALDAIIASVTESLKEGDD<br>ALDAIIASVTESLKEGDDVAL | 3 | 0 | 24-40<br>24-41<br>24-44 | 12 | ALDAIIASVTESLKE<br>ALDAIIASVTESLKEGDDVALVG | 2 | 1 | 24-38<br>24-46 | 7 |
| P0ACF8 | REEESAAAAEVE<br>REEESAAAAEVEE | 2 | 1 | 41-52<br>41-53 | 5 | REEESAAAAE<br>REEESAAAAEVE | 2 | 1 | 41-50<br>41-52 | 9 | REEESAAAAEVE<br>REEESAAAAEVEE | 2 | 2 | 41-52<br>41-53 | 11 | REEESAAAAE<br>REEESAAAAEV<br>REEESAAAAEVE<br>REEESAAAAEVEE | 4 | 3 | 41-50<br>41-51<br>41-52<br>41-53 | 26 |
| P0AD33 | ITPTFTEESDGVRL<br>ITPTFTEESDGVRLD<br>ITPTFTEESDGVRLDID | 3 | 1 | 61-74<br>61-75<br>61-77 | 10 | | | | | | | | | | | | | | | |

(Supplementary Table S1, continued)

| UniProt ID | No treatment | | | | | Chloramphenicol | | | | | Kasugamycin & Puromycin | | | | | Erythromycin | | | | |
|---|---|---|---|---|---|---|---|---|---|---|---|---|---|---|---|---|---|---|---|---|
| | All_seq | Number of Detected Peptides | Number of Detected Tris-added peptides | Detected Position | Sum of corre-sponding PSM numbers | All_seq | Number of Detected Peptides | Number of Detected Tris-added peptides | Detected Position | Sum of corre-sponding PSM numbers | All_seq | Number of Detected Peptides | Number of Detected Tris-added peptides | Detected Position | Sum of corre-sponding PSM numbers | All_seq | Number of Detected Peptides | Number of Detected Tris-added peptides | Detected Position | Sum of corre-sponding PSM numbers |
| P0ADZ4 | | | | | | | | | | | | | | | | DANDTGSTEVQVALLTAQINH<br>DANDTGSTEVQVALLTAQINHL<br>DANDTGSTEVQVALLTAQINHLQGH<br>DANDTGSTEVQVALLTAQINHLQGHFAE | 4 | 0 | 18-38<br>18-39<br>18-42<br>18-45 | 18 |
| P0AES9 | | | | | | | | | | | | | | | | KPVNSWTCEDFLAVD<br>KPVNSWTCEDFLAVDES | 2 | 1 | 32-46<br>32-48 | 7 |
| P0AFL3 | | | | | | | | | | | | | | | | APVSVQNFVDY<br>APVSVQNFVDYVNS<br>APVSVQNFVDYVNSGFYNNTTF | 3 | 0 | 49-59<br>49-62<br>49-70 | 8 |
| P0AFV4 | | | | | | | | | | | | | | | | GIPAIAVAVLLSAC<br>GIPAIAVAVLLSACSANN | 2 | 0 | 14-27<br>14-31 | 31 |
| P0AGB3 | | | | | | | | | | | | | | | | NYAGYGLPQADL<br>NYAGYGLPQADLIQE | 2 | 0 | 67-78<br>67-81 | 5 |
| P13445 | | | | | | VLDATQLYLGEIGY<br>VLDATQLYLGEIGYSPLLTAE | 2 | 0 | 54-67<br>54-74 | 8 | | | | | | | | | | |
| P62399 | | | | | | KLLDNAAA<br>KLLDNAAADLAAISG<br>KLLDNAAADLAAISGQ | 3 | 2 | 48-55<br>48-62<br>48-63 | 18 | | | | | | | | | | |
| | | | | | | LLDNAAADLAAISG<br>LLDNAAADLAAISGQ | 2 | 1 | 49-62<br>49-63 | 14 | | | | | | | | | | |
| P67910 | | | | | | FVNLVDLNIAD<br>FVNLVDLNIADYMDKED | 2 | 0 | 39-49<br>39-55 | 8 | | | | | | FVNLVDLNIADY<br>FVNLVDLNIADYMDKEDF | 2 | 0 | 39-50<br>39-56 | 5 |
| P69776 | LVLGAVILGSTLLAG<br>LVLGAVILGSTLLAGC<br>LVLGAVILGSTLLAGCS<br>LVLGAVILGSTLLAGCSS<br>LVLGAVILGSTLLAGCSSN | 5 | 3 | 6-20<br>6-21<br>6-22<br>6-23<br>6-24 | 34 | LVLGAVILGSTL<br>LVLGAVILGSTLLAG<br>LVLGAVILGSTLLAGC<br>LVLGAVILGSTLLAGCS<br>LVLGAVILGSTLLAGCSS<br>LVLGAVILGSTLLAGCSSN | 6 | 2 | 6-17<br>6-20<br>6-21<br>6-22<br>6-23<br>6-24 | 72 | LVLGAVILGSTLLAG<br>LVLGAVILGSTLLAGC<br>LVLGAVILGSTLLAGCSSN | 3 | 0 | 6-20<br>6-21<br>6-24 | 11 | LVLGAVILGSTLLAG<br>LVLGAVILGSTLLAGC<br>LVLGAVILGSTLLAGCS<br>LVLGAVILGSTLLAGCSS<br>LVLGAVILGSTLLAGCSSN | 5 | 2 | 6-20<br>6-21<br>6-22<br>6-23<br>6-24 | 118 |
| P69831 | | | | | | NFFELEGIAIPHGTSA<br>NFFELEGIAIPHGTSAY | 2 | 0 | 170-185<br>170-186 | 4 | | | | | | | | | | |
| P77454 | | | | | | | | | | | | | | | | IGADPTGLPFNSVIAL<br>IGADPTGLPFNSVIALELH<br>IGADPTGLPFNSVIALELHGGKPL<br>IGADPTGLPFNSVIALELHGGKPLSPL | 4 | 0 | 89-104<br>89-107<br>89-112<br>89-115 | 9 |
| P77747 | | | | | | VDGLHYFSD<br>VDGLHYFSDNKDVDGD<br>VDGLHYFSDNKDVDGDQTY | 3 | 1 | 38-46<br>38-53<br>38-56 | 20 | | | | | | | | | | |

**Table S2. Preparation of the templated DNA fragments used for in vitro translation.**

| feature | 1st PCR | | | 2nd PCR | | | Experiment |
|---|---|---|---|---|---|---|---|
| | template DNA | primer_1 | primer_2 | template DNA | primer_1 | primer_2 | |
| folA_N10_nonstop | pET-DHFR | CGCGAAATTAATACGACTCACTATAGGG | TACCGCTAACGCCGCAATCAGAC | | | | Fig. 1B |
| folA_N20_nonstop | pET-DHFR | CGCGAAATTAATACGACTCACTATAGGG | CATGGCGTTTTCCATGCCGATAAC | | | | Fig. 1B |
| folA_N40_nonstop | pET-DHFR | CGCGAAATTAATACGACTCACTATAGGG | CACGGGTTTATTTAAGGTGTTG | | | | Fig. 1B |
| folA_N80_nonstop | pET-DHFR | CGCGAAATTAATACGACTCACTATAGGG | TTCATCCACCGACTTCACCCACG | | | | Fig. 1B |
| folA_N159+Ala_nonstop | pET-DHFR | CGCGAAATTAATACGACTCACTATAGGG | TGCCCGCCGCTCCAGAATCTCAAAG | | | | Fig. 1B |
| eno_N10_nonstop | pET-ENO | CGCGAAATTAATACGACTCACTATAGGG | ACGACCGATGATTTTTACGATTTTGG | | | | Fig. 1B |
| eno_N20_nonstop | pET-ENO | CGCGAAATTAATACGACTCACTATAGGG | AGTCGGGTTACCACGGGAGTCG | | | | Fig. 1B |
| eno_N40_nonstop | pET-ENO | CGCGAAATTAATACGACTCACTATAGGG | ACCTGACGGAGCAGCTGCCATACC | | | | Fig. 1B |
| eno_N80_nonstop | pET-ENO | CGCGAAATTAATACGACTCACTATAGGG | AATCAGCGCCTGAGCGATCGGG | | | | Fig. 1B |
| eno_N160_nonstop | pET-ENO | CGCGAAATTAATACGACTCACTATAGGG | AGCGTGCTCACCACCGTTGATG | | | | Fig. 1B |
| sfGFP_N10_nonstop | pET-sfGFP | CGCGAAATTAATACGACTCACTATAGGG | TCCAGTGAAAAGTTCTTCTCC | | | | Fig. 1B |
| sfGFP_N20_nonstop | pET-sfGFP | CGCGAAATTAATACGACTCACTATAGGG | ACCATCTAATTCAACAAGAATTG | | | | Fig. 1B |
| sfGFP_N40_nonstop | pET-sfGFP | CGCGAAATTAATACGACTCACTATAGGG | TCCGTTTGTTGCATCACCTTCACC | | | | Fig. 1B |
| sfGFP_N80_nonstop | pET-sfGFP | CGCGAAATTAATACGACTCACTATAGGG | CCGTTTCATGTGATCCGGATAACG | | | | Fig. 1B |
| sfGFP_N160_nonstop | pET-sfGFP | CGCGAAATTAATACGACTCACTATAGGG | TCCATTCTTTTGTTTGTCTGCC | | | | Fig. 1B |
| secM_N10_nonstop | pCA24N-secM | TTTAACTTTAAGAAGGAGATATACCAATGAGTGGAATACTGACGCGCTG | CTGTCGCCAGCGCGTCAGTATTCCAC | 1st PCR | GAAATTAATACGACTCACTATAGGGAGACCACAACGGTTTCCCTCTAGAAATAATTTTGTTTAACTTTAAGAAGGAG | CTGTCGCCAGCGCGTCAGTATTCCAC | Fig. 1B, Fig. S1B |
| secM_N20_nonstop | pCA24N-secM | TTTAACTTTAAGAAGGAGATATACCAATGAGTGGAATACTGACGCGCTG | GAGATGCGGCCAGAAGTAGCGTTTAC | 1st PCR | GAAATTAATACGACTCACTATAGGGAGACCACAACGGTTTCCCTCTAGAAATAATTTTGTTTAACTTTAAGAAGGAG | GAGATGCGGCCAGAAGTAGCGTTTAC | Fig. 1B, Fig. S1B |
| secM_N40_nonstop | pCA24N-secM | TTTAACTTTAAGAAGGAGATATACCAATGAGTGGAATACTGACGCGCTG | TGGTTCGGCGGCGTTGCTGAGCGCAG | 1st PCR | GAAATTAATACGACTCACTATAGGGAGACCACAACGGTTTCCCTCTAGAAATAATTTTGTTTAACTTTAAGAAGGAG | TGGTTCGGCGGCGTTGCTGAGCGCAG | Fig. 1B, Fig. S1B |
| secM_N80_nonstop | pCA24N-secM | TTTAACTTTAAGAAGGAGATATACCAATGAGTGGAATACTGACGCGCTG | GTAATCAACGGAATAGTTCGAATTCGGG | 1st PCR | GAAATTAATACGACTCACTATAGGGAGACCACAACGGTTTCCCTCTAGAAATAATTTTGTTTAACTTTAAGAAGGAG | GTAATCAACGGAATAGTTCGAATTCGGG | Fig. 1B, Fig. S1B |
| secM_N160_nonstop | pCA24N-secM | TTTAACTTTAAGAAGGAGATATACCAATGAGTGGAATACTGACGCGCTG | TTGCGCCTGGCTTATCCAGACGGGCGTGCTG | 1st PCR | GAAATTAATACGACTCACTATAGGGAGACCACAACGGTTTCCCTCTAGAAATAATTTTGTTTAACTTTAAGAAGGAG | TTGCGCCTGGCTTATCCAGACGGGCGTGCTG | Fig. 1B, Fig. S1B |
| sfGFP_N40_10A_nonstop | pET-sfGFP | CGCGAAATTAATACGACTCACTATAGGG | AGCAGCAGCAGCAGCAGCAGCAGCAGCAGCTCCGTTTGTTGCATCACCTTCACC | | | | Fig. S1C |
| sfGFP_N40_10C_nonstop | pET-sfGFP | CGCGAAATTAATACGACTCACTATAGGG | GCAGCAGCAGCAGCAGCAGCAGCAGCAGCATCCGTTTGTTGCATCACCTTCACC | | | | Fig. S1C |
| sfGFP_N40_10D_nonstop | pET-sfGFP | CGCGAAATTAATACGACTCACTATAGGG | GTCGTCGTCGTCGTCGTCGTCGTCGTCGTCTCCGTTTGTTGCATCACCTTCACC | | | | Fig. S1C |
| sfGFP_N40_10E_nonstop | pET-sfGFP | CGCGAAATTAATACGACTCACTATAGGG | TTCTTCTTCTTCTTCTTCTTCTTCTTCTTCTCCGTTTGTTGCATCACCTTCACC | | | | Fig. S1C |
| sfGFP_N40_10F_nonstop | pET-sfGFP | CGCGAAATTAATACGACTCACTATAGGG | GAAGAAGAAGAAGAAGAAGAAGAAGAAGAATCCGTTTGTTGCATCACCTTCACC | | | | Fig. S1C |
| sfGFP_N40_10G_nonstop | pET-sfGFP | CGCGAAATTAATACGACTCACTATAGGG | ACCACCACCACCACCACCACCACCACCACCTCCGTTTGTTGCATCACCTTCACC | | | | Fig. S1C |
| sfGFP_N40_10H_nonstop | pET-sfGFP | CGCGAAATTAATACGACTCACTATAGGG | GTGGTGGTGGTGGTGGTGGTGGTGGTGGTGTCCGTTTGTTGCATCACCTTCACC | | | | Fig. S1C |
| sfGFP_N40_10I_nonstop | pET-sfGFP | CGCGAAATTAATACGACTCACTATAGGG | GATGATGATGATGATGATGATGATGATGATTCCGTTTGTTGCATCACCTTCACC | | | | Fig. S1C |
| sfGFP_N40_10K_nonstop | pET-sfGFP | CGCGAAATTAATACGACTCACTATAGGG | CTTCTTCTTCTTCTTCTTCTTCTTCTTCTTTCCGTTTGTTGCATCACCTTCACC | | | | Fig. S1C |
| sfGFP_N40_10L_nonstop | pET-sfGFP | CGCGAAATTAATACGACTCACTATAGGG | CAGCAGCAGCAGCAGCAGCAGCAGCAGCAGTCCGTTTGTTGCATCACCTTCACC | | | | Fig. S1C |
| sfGFP_N40_10M_nonstop | pET-sfGFP | CGCGAAATTAATACGACTCACTATAGGG | CATCATCATCATCATCATCATCATCATCATTCCGTTTGTTGCATCACCTTCACC | | | | Fig. S1C |
| sfGFP_N40_10N_nonstop | pET-sfGFP | CGCGAAATTAATACGACTCACTATAGGG | GTTGTTGTTGTTGTTGTTGTTGTTGTTGTTTCCGTTTGTTGCATCACCTTCACC | | | | Fig. S1C |
| sfGFP_N40_10P_nonstop | pET-sfGFP | CGCGAAATTAATACGACTCACTATAGGG | TGGTGGTGGTGGTGGTGGTGGTGGTGGTGGTCCGTTTGTTGCATCACCTTCACC | | | | Fig. S1C |
| sfGFP_N40_10Q_nonstop | pET-sfGFP | CGCGAAATTAATACGACTCACTATAGGG | CTGCTGCTGCTGCTGCTGCTGCTGCTGCTGTCCGTTTGTTGCATCACCTTCACC | | | | Fig. S1C |
| sfGFP_N40_10R_nonstop | pET-sfGFP | CGCGAAATTAATACGACTCACTATAGGG | ACGACGACGACGACGACGACGACGACGACGTCCGTTTGTTGCATCACCTTCACC | | | | Fig. S1C |
| sfGFP_N40_10S_nonstop | pET-sfGFP | CGCGAAATTAATACGACTCACTATAGGG | GCTGCTGCTGCTGCTGCTGCTGCTGCTGCTTCCGTTTGTTGCATCACCTTCACC | | | | Fig. S1C |
| sfGFP_N40_10T_nonstop | pET-sfGFP | CGCGAAATTAATACGACTCACTATAGGG | GGTGGTGGTGGTGGTGGTGGTGGTGGTGGTTCCGTTTGTTGCATCACCTTCACC | | | | Fig. S1C |
| sfGFP_N40_10V_nonstop | pET-sfGFP | CGCGAAATTAATACGACTCACTATAGGG | TACTACTACTACTACTACTACTACTACTACTCCGTTTGTTGCATCACCTTCACC | | | | Fig. S1C |
| sfGFP_N40_10W_nonstop | pET-sfGFP | CGCGAAATTAATACGACTCACTATAGGG | CCACCACCACCACCACCACCACCACCACCATCCGTTTGTTGCATCACCTTCACC | | | | Fig. S1C |
| sfGFP_N40_10Y_nonstop | pET-sfGFP | CGCGAAATTAATACGACTCACTATAGGG | GTAGTAGTAGTAGTAGTAGTAGTAGTAGTATCCGTTTGTTGCATCACCTTCACC | | | | Fig. S1C |

Supplementary Table S3. Settings of Proteome Discoverer 2.4.

Processing Workflow:

| Node | Parameter | Values |
|---|---|---|
| Spectrum Files | (No parameters) | |
| Spectrum Selector | 1. General Settings | |
| | Precursor Selection | Use MS1 Precursor |
| | Use Isotope Pattern in Precursor Reevaluation | True |
| | Provide Profile Spectra | Automatic |
| | 2. Spectrum Properties Filter | |
| | Lower RT Limit | 0 |
| | Upper RT Limit | 0 |
| | First Scan | 0 |
| | Last Scan | 0 |
| | Lowest Charge State | 0 |
| | Highest Charge State | 0 |
| | Min. Precursor Mass | 350 Da |
| | Max. Precursor Mass | 5000 Da |
| | Total Intensity Threshold | 0 |
| | Minimum Peak Count | 1 |
| | 3. Scan Event Filters | |
| | MS Order | Is Not MS1 |
| | Min. Collision Energy | 0 |
| | Max. Collision Energy | 1000 |
| | Scan Type | Is Full |
| | 4. Peak Filters | |
| | S/N Threshold (FT-only) | 1.5 |
| | 5. Replacements for Unrecognized Properties | |
| | Unrecognized Charge Replacements | Automatic |
| | Unrecognized Mass Analyzer Replacements | ITMS |
| | Unrecognized MS Order Replacements | MS2 |
| | Unrecognized Activation Type Replacements | CID |
| | Unrecognized Polarity Replacements | + |
| | Unrecognized MS Resolution@200 Replacements | 60000 |
| | Unrecognized MSn Resolution@200 Replacements | 30000 |
| | 6. Precursor Pattern Extraction | |
| | Precursor Clipping Range Before | 2.5 Da |
| | Precursor Clipping Range After | 5.5 Da |
| Sequest HT | 1. Input Data | |
| | Protein Database | PD_Contaminants_2015_5 (bundled in Proteome Discoverer)<br>All *E.coli* ORFs (from UniProt: UP000000625, downloaded on 190820) |
| | Enzyme | Trypsin (Semi) |
| | Max. Missed Cleavage Sites | 2 |
| | Min. Peptide Length | 6 |
| | Max. Peptide Length | 100 |
| | Max. Number of Peptides Reported | 10 |

| | | |
|---|---|---|
| | 2. Tolerances | |
| | Precursor Mass Tolerance | 10 ppm |
| | Fragment Mass Tolerance | 0.02 Da |
| | Use Average Precursor Mass | False |
| | Use Average Fragment Mass | False |
| | 3. Spectrum Matching | |
| | Use Neutral Loss a Ions | True |
| | Use Neutral Loss b Ions | True |
| | Use Neutral Loss y Ions | True |
| | Use Flanking Ions | True |
| | Weight of a Ions | 0 |
| | Weight of b Ions | 1 |
| | Weight of c Ions | 0 |
| | Weight of x Ions | 0 |
| | Weight of y Ions | 1 |
| | Weight of z Ions | 0 |
| | 3. Spectrum Matching | |
| | Use Neutral Loss a Ions | True |
| | Use Neutral Loss b Ions | True |
| | Use Neutral Loss y Ions | True |
| | Use Flanking Ions | True |
| | Weight of a Ions | 0 |
| | Weight of b Ions | 1 |
| | Weight of c Ions | 0 |
| | Weight of x Ions | 0 |
| | Weight of y Ions | 1 |
| | Weight of z Ions | 0 |
| | 4. Dynamic Modifications | |
| | Max. Equal Modifications Per Peptide | 3 |
| | Max. Dynamic Modifications Per Peptide | 4 |
| | Dynamic Modification | Oxidation / +15.995 Da (M)<br>Deamidated / +0.984 Da (N, Q) |
| | 5. Dynamic Modifications (peptide terminus) | |
| | C-Terminal Modification | Tris in Cterm / +103.063 Da (C-Terminus) |
| | 6. Dynamic Modifications (protein terminus) | |
| | 1. N-Terminal Modification | Acetyl / +42.011 Da (N-Terminus) |
| | 7. Static Modifications | |
| | 1. Static Modification | Carbamidomethyl / +57.021 Da (C) |
| Percolator | 1. Target/Decoy Strategy | |
| | Target/Decoy Selection | Concatenated |
| | Validation based on | q-Value |
| | 2. Input Data | |
| | Maximum Delta Cn | 0.05 |
| | Maximum Rank | 0 |
| | 3. FDR Targets | |
| | Target FDR (Strict) | 0.01 |
| | Target FDR (Relaxed) | 0.05 |
| Minora Feature Detector | 1. Peak & Feature Detection | |
| | Min. Trace Length | 5 |
| | Max. deltaRT of Isotope Pattern Multiplets [min] | 0.2 |
| | 2. Feature to ID Linking | |
| | PSM Confidence At Least | High |

Consensus Workflow:

| Node | Parameter | Values |
|---|---|---|
| MSF Files | 1. Storage Settings | |
| | Spectra to Store | Identified or Quantified |
| | Feature Traces to Store | All |
| | 2. Merging of Identified Peptide and Proteins | |
| | Merge Mode | Do Not Merge |
| | 3. FASTA Title Line Display | |
| | Reported FASTA Title Lines | Best match |
| | Title Line Rule | standard |
| | 4. PSM Filters | |
| | Maximum Delta Cn | 0.05 |
| | Maximum Rank | 0 |
| | Maximum Delta Mass | 0 ppm |
| PSM Grouper | 1. Peptide Group Modifications | |
| | Site Probability Threshold | 75 |
| Peptide Validator | 1. General Validation Settings | |
| | Validation Mode | Automatic (Control peptide level error rate if possible) |
| | Target FDR (Strict) for PSMs | 0.01 |
| | Target FDR (Relaxed) for PSMs | 0.05 |
| | Target FDR (Strict) for Peptides | 0.01 |
| | 2. Specific Validation Settings | |
| | Validation Based on | q-Value |
| | Target/Decoy Selection for PSM Level FDR Calculation Based on Score | Automatic |
| | Reset Confidences for Nodes without Decoy Search (Fixed score thresholds) | False |
| Peptide and Protein Filter | 1. Peptide Filters | |
| | Peptide Confidence At Least | High |
| | Keep Lower Confident PSMs | True |
| | Minimum Peptide Length | 6 |
| | Remove Peptides Without Protein Reference | False |
| | 2. Protein Filters | |
| | Minimum Number of Peptide Sequences | 1 |
| | Count Only Rank 1 Peptides | False |
| | Count Peptides Only for Top Scored Protein | False |
| Protein Scorer | (No parameters) | |
| Protein Grouping | 1. Protein Grouping | |
| | Apply strict parsimony principle | True |
| Peptide in Protein Annotation | 1. Flanking Residues | |
| | Annotate Flanking Residues of the Peptide | True |
| | Number Flanking Residues in Connection Tables | 1 |
| | 2. Modifications in Peptide | |
| | Protein Modifications Reported | Only for Master Proteins |
| | 3. Modifications in Protein | |
| | Modification Sites Reported | All And Specific |

| | | |
|---|---|---|
| | Minimum PSM Confidence | High |
| | Report Only PTMs | True |
| | 4. Positions in Protein | |
| | Protein Positions for Peptides | Only for Master Proteins |
| Protein FDR Validator | 1. Confidence Thresholds | |
| | Target FDR (Strict) | 0.01 |
| | Target FDR (Relaxed) | 0.05 |
| Protein Marker | 1. Contaminant Database | |
| | Protein Database | PD_Contaminants_2015_5 |
| | 5. Annotate Species | |
| | As Species Map | False |
| | As Species Names | False |
| | 6. Mark Additional Entities | |
| | Annotation Groups | False |
| | Pathway Groups | False |
| | Modification Sites | True |
| | Peptide Isoform Groups | True |
| Feature Mapper | 1. Chromatographic Alignment | |
| | Perform RT Alignment | True |
| | Maximum RT Shift [min] | 10 |
| | Mass Tolerance | 10 ppm |
| | Parameter Tuning | Coarse |
| | 2. Feature Linking and Mapping | |
| | RT Tolerance [min] | 0 |
| | Mass Tolerance | 0 ppm |
| | Min. S/N Threshold | 5 |
| Precursor Ions Quantifier (only for quantification) | 1. General Quantification Settings | |
| | Peptides to Use | Unique + Razor |
| | Consider Protein Groups for Peptide Uniqueness | True |
| | Use Shared Quan Results | True |
| | Reject Quan Results with Missing Channels | False |
| | 2. Precursor Quantification | |
| | Precursor Abundance Based On | Intensity |
| | Min. # Replicate Features [%] | 0 |
| | 3. Normalization and Scaling | |
| | Normalization Mode | Total Peptide Amount |
| | Scaling Mode | None |
| | 4. Exclude Peptides from Protein Quantification | |
| | For Normalization | Use All Peptides |
| | For Protein Roll-Up | Use All Peptides |
| | For Pairwise Ratios | Exclude Modified |
| | 5. Quan Rollup and Hypothesis Testing | |
| | Protein Abundance Calculation | Summed Abundances |
| | N for Top N | 3 |
| | Protein Ratio Calculation | Protein Abundance Based |
| | Maximum Allowed Fold Change | 1000 |
| | Imputation Mode | None |
| | Hypothesis Test | ANOVA (Individual Proteins) |
| | 6. Quan Ratio Distributions | |
| | 1st Fold Change Threshold | 2 |
| | 2nd Fold Change Threshold | 4 |
| | 3rd Fold Change Threshold | 6 |
| | 4th Fold Change Threshold | 8 |
| | 5th Fold Change Threshold | 10 |
